# Snapshots of *Pseudomonas aeruginosa* SOS response activation complex reveal structural prerequisites for LexA engagement and cleavage

**DOI:** 10.1101/2024.03.22.585941

**Authors:** Filippo Vascon, Sofia De Felice, Matteo Gasparotto, Stefan T. Huber, Claudio Catalano, Monica Chinellato, Alessandro Grinzato, Francesco Filippini, Lorenzo Maso, Arjen J. Jakobi, Laura Cendron

## Abstract

Antimicrobial resistance represents a major threat to human health and *Pseudomonas aeruginosa* stands out among the pathogens responsible for this emergency. The SOS response to DNA damage plays a pivotal role in bacterial evolution, driving the development of resistance mechanisms and influencing the adaptability of bacterial populations to challenging environments, particularly in the context of antibiotic exposure. Recombinase A (RecA) and the transcriptional repressor LexA are the key players that orchestrate this process, determining either the silencing or the active transcription of the genes under their control. By integrating state-of-the-art structural approaches with binding and functional assays *in vitro*, we elucidated the molecular events governing the SOS response activation in *P. aeruginosa*, focusing on the RecA-LexA interaction. Our findings identify the conserved determinants and strength of the interactions that let RecA trigger the autocleavage and inactivation of the LexA repressor. These results provide the groundwork for designing novel antimicrobial strategies and for exploring the potential translation of *Escherichia coli*-derived approaches, to address the health-threatening implications of bacterial infections.

**Graphical Abstract:** 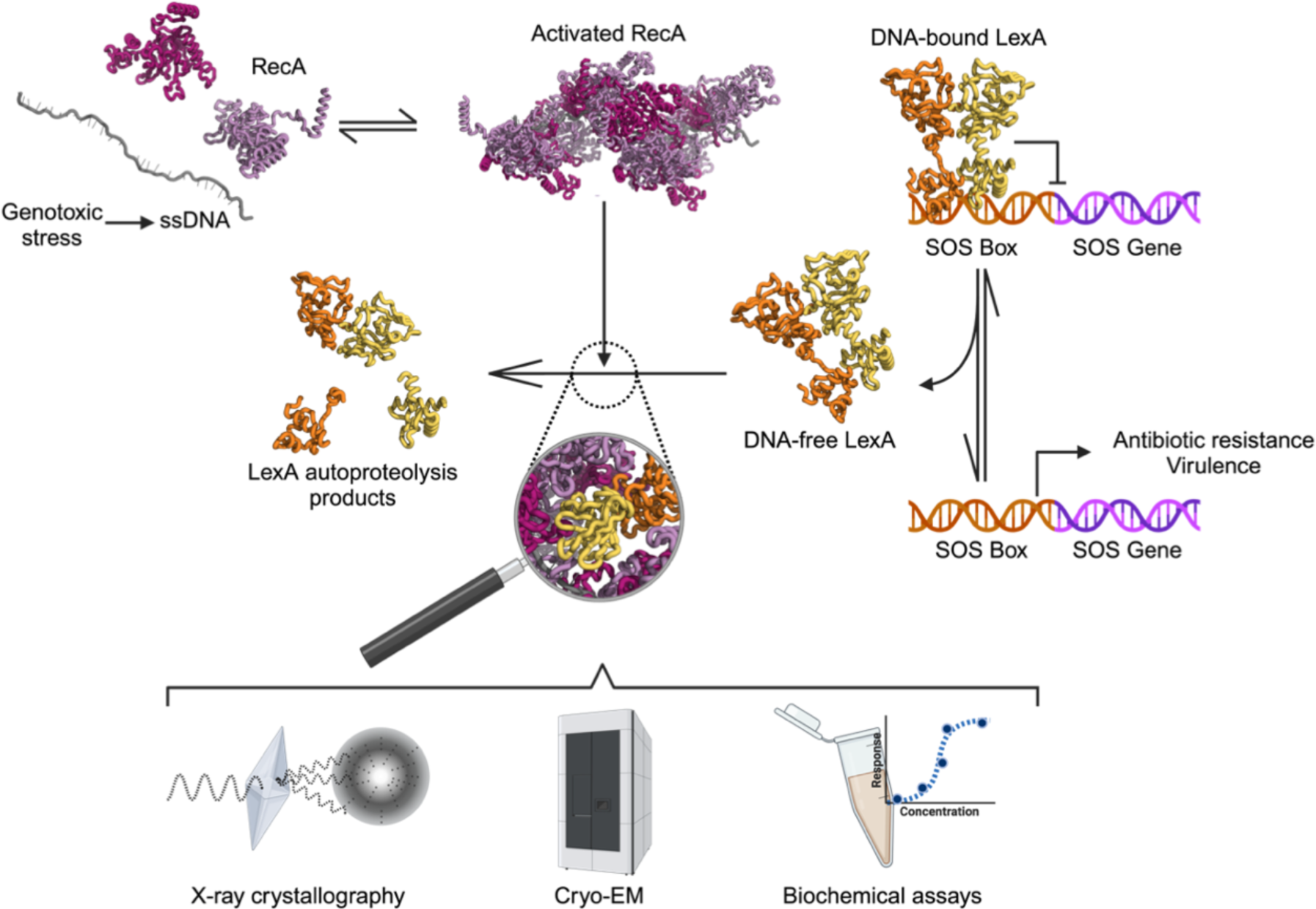

## Introduction

To guide and coordinate the development of novel antimicrobial strategies, several national and international health agencies constantly monitor the prevalence of antibiotic resistant bacterial pathogens, prioritizing those representing the greatest threats (CDC, 2019; Tacconelli *et al*, 2018). The Gram-negative bacterium *Pseudomonas aeruginosa* always finds its spot in these “priority lists”, as it displays a vast spectrum of antibiotic resistance mechanisms (Pang *et al*, 2019) and a high frequency of infections among hospitalized patients, either as a direct etiologic agent or as a comorbidity, occasionally acquired in the healthcare settings (Rice, 2008). Indeed, as an opportunistic pathogen, *P. aeruginosa* mainly infects patients suffering from immune deficiencies, severe wounds, and pulmonary diseases, including cystic fibrosis and COVID-19 (Liao *et al*, 2022; Qin *et al*, 2022).

Together with multi-drug resistance, a notable variety of virulence factors determines *P. aeruginosa* pathogenicity and recalcitrance. Several surface appendages (pili) and proteins (e.g., lectins) mediate *P. aeruginosa* adhesion to the host tissues (Liao *et al*, 2022; Pang *et al*, 2019), while secreted proteases and toxins damage the host’s tissue components, immune defenses and physiological functions (Liao *et al*, 2022; Qin *et al*, 2022). *P. aeruginosa* is known to form biofilms and communicate via quorum sensing (QS). These interwoven features are of great relevance in the fight against bacterial pathogens, since biofilms physically shield the enclosed sensitive cells from the action of antimicrobials and favor the differentiation of persister sub-populations (Pang *et al*, 2019; Podlesek & Žgur Bertok, 2020), while QS regulates the expression of virulence factors (Qin *et al*, 2022).

In recent years, anti-evolutive, anti-virulence, anti-biofilm, and quorum quencher strategies have been proposed as new approaches in antimicrobial chemotherapy, as they could counteract the rapid acquisition of antibiotic resistance and weaken the pathogenicity of bacterial infections (Mulani *et al*, 2019; Merrikh & Kohli, 2020; Culyba *et al*, 2015; Mühlen & Dersch, 2016).

A master regulator involved in the control of cell division, fitness to environmental stressors, prophage activation, biofilm maturation, production of virulence factors, and error-prone DNA replication is represented by the SOS response pathway (Gotoh *et al*, 2010; Lima-Noronha *et al*, 2022; Galhardo *et al*, 2009; Pacheco & Sperandio, 2012). Most importantly, it is the most conserved mechanism of bacterial response to DNA damage induced by the exposure to antimicrobials, UV radiation, and reactive oxygen species. Because of these reasons, it is regarded as one of the best targets of anti-evolutive and antivirulence therapies (Culyba *et al*, 2015; Merrikh & Kohli, 2020; Dawan & Ahn, 2022).

The plethora of SOS-regulated mechanisms is species-specific and depends on the set of genes (the *SOS regulon*) controlled by the master SOS transcriptional repressor LexA through its binding to specific operator sequences in the promoter region of SOS genes (*SOS boxes;* Zhang *et al*, 2010).

A prerequisite for triggering the SOS response is the activation of Recombinase A (RecA), which senses single-stranded DNA generated by the genotoxic damage and oligomerizes on it in an ATP-dependent manner. RecA oligomers promote the autoproteolytic cleavage of the dimeric LexA, in its DNA-free form (Butala *et al*, 2011). This activity is exerted by a Ser/Lys dyad (S125/K162 in *P. aeruginosa*) on a *scissile* peptide bond (A90-G91 in *P. aeruginosa*) located on a flexible loop, which can switch between an inactive (open) and a prone-to-cleavage (closed) conformation (Luo *et al*, 2001; Mo *et al*, 2014).

The autoproteolysis event hinders the transcriptional repressor activity of the cleavage products (*i.e*. LexA N-terminal and C-terminal domains, NTD and CTD) and shifts the equilibrium between the DNA-bound and unbound LexA towards the latter state. LexA autoproteolysis thus leads to the active expression of the SOS genes, with tightly regulated expression levels, chronological order, and duration, that depend on LexA affinity and binding kinetics on the different SOS boxes (Culyba *et al*, 2018; Zhang *et al*, 2010).

Despite the species-specificity of the SOS regulon – e.g. it accounts for 57 genes in *E. coli* (Simmons *et al*, 2008), 33 genes in *Bacillus subtilis,* 48 genes in *Salmonella enterica* (Mérida-Floriano *et al*, 2021) and 15 genes in *P. aeruginosa* (Cirz *et al*, 2006) – it invariably includes factors involved in DNA repair, in particular error-prone translesion (TLS) DNA polymerases (Erill *et al*, 2007). Despite less studied than the SOS-regulated *Pol II, Pol IV* and *Pol V* of *E. coli*, other error-prone DNA polymerases (ImuB and ImuC, also known as DnaE2) encoded by SOS-inducible *imuA-imuB-dnaE2* gene cassettes are broadly distributed among bacterial taxa, including *P. aeruginosa* (Jatsenko *et al*, 2017; Luján *et al*, 2019; Erill *et al*, 2007), confirming the centrality of translesion synthesis in the general SOS response. These TLS polymerases can bypass DNA lesions otherwise incompatible with replicative polymerases, at the cost of high error rates, thus introducing mutations (Fujii & Fuchs, 2020). As a result, one of the primary outcomes of the SOS response is a transient hypermutator state that promotes genetic diversity, adaptive mutation and the evolution of antimicrobial resistance. Given its importance for the acquisition of antimicrobial resistance and its high conservation, the SOS response is currently receiving attention as a target of antibiotic-adjunctive therapies, that might prolong antibiotics effectiveness and even increase their efficacy (Lu & Collins, 2009; Maso *et al*, 2022; Mo *et al*, 2018; Bellio *et al*, 2017; Yakimov *et al*, 2017; Barreto *et al*, 2009; Selwood *et al*, 2018).

While the structural features of the single components LexA and RecA have been determined by X-ray crystallography or cryogenic electron microscopy (Cryo-EM), a substantial lack of structural and mechanistic knowledge about the SOS complex has limited our comprehension of the stimulatory role played by RecA toward LexA autocleavage. Only recently, Cryo-EM studies on the SOS complex of *E. coli* began to shed light on the interaction site of either LexA C-terminal domain or full-length protein with RecA/ssDNA/ATPγS oligomers (Gao *et al*, 2023; preprint: Cory *et al*, 2023).

The cascade of events promoted by DNA damage in *P. aeruginosa* (Pa) still needs a complete characterization, and several recent works have unveiled a previously unknown complexity compared to the well-studied *E. coli* (Ec) model (e.g., multiple LexA-like transcriptional regulators and interconnections with other stress-response pathways; Penterman *et al*, 2014; Jiao *et al*, 2021; Fan *et al*, 2019). Deepening our understanding of the principal protein actors of *P. aeruginosa* SOS response is necessary to determine to which extent the approaches developed in *E. coli* could be translated to this pathogen.

With this aim, our work investigated the core of the SOS response in *P. aeruginosa*, obtaining the structures of the isolated components (LexA_Pa_ C-terminal domain and RecA_Pa_/ssDNA/ATPγS), as well as the Cryo-EM structure of the activation complex (LexA_Pa_S125A-RecA_Pa_/ssDNA/ATPγS assembly). Our structural data, integrated by experimental measurements of the affinity of the binding partners and proteolysis assays, let us describe the molecular events governing the binding and activation of the SOS response players in this health-threatening pathogen.

## Results

### Crystal structure of LexA_Pa_^CTD^ G91D

Two mutants of *P. aeruginosa* LexA were expressed in *E. coli*, purified by affinity chromatography, and used for the structural studies described in this work, which require a stable LexA variant unable to undergo RecA*-dependent or independent autoproteolysis. Specifically, the LexA_Pa_S125A mutant consists of the full-length protein carrying the S125A mutation in the catalytic dyad. Conversely, LexA_Pa_^CTD^ G91D comprises only the LexA_Pa_ C-terminal domain (CTD, from Gly81 to Arg204) bearing an inactivating mutation on the cleavage site. While the former will be used to study RecA_Pa_-LexA_Pa_ interaction (as reported below), the latter is more amenable to protein crystallization as it lacks the flexible linker and NTD.

In agreement with previous observations (Mohana-Borges *et al*, 2000), analytical size exclusion chromatography showed that both proteins behave as homodimers in solution (Fig. 1 A). Specifically, LexA_Pa_S125A was eluted with an apparent molecular weight of 62 ± 6 kDa, and LexA_Pa_^CTD^ G91D eluted at an apparent molecular weight of 35 ± 4 kDa, in both cases corresponding to roughly the double of the expected molecular weight of the monomeric forms (24 kDa and 14 kDa, respectively). Moreover, SDS-PAGE-based analysis of RecA_Pa_/ssDNA/ATPγS (RecA_Pa_*)-induced autoproteolysis reactions revealed similar self-cleavage kinetics for wild-type full-length LexA_Pa_ and LexA_Pa_^CTD^, while both the S125A and G91D mutations completely abated the catalytic activity of the LexA_Pa_ variants (Fig. 1 B). These observations confirmed that the C-terminal domain provides all the determinants for LexA homodimerization and autoproteolysis.

**Fig. 1:**
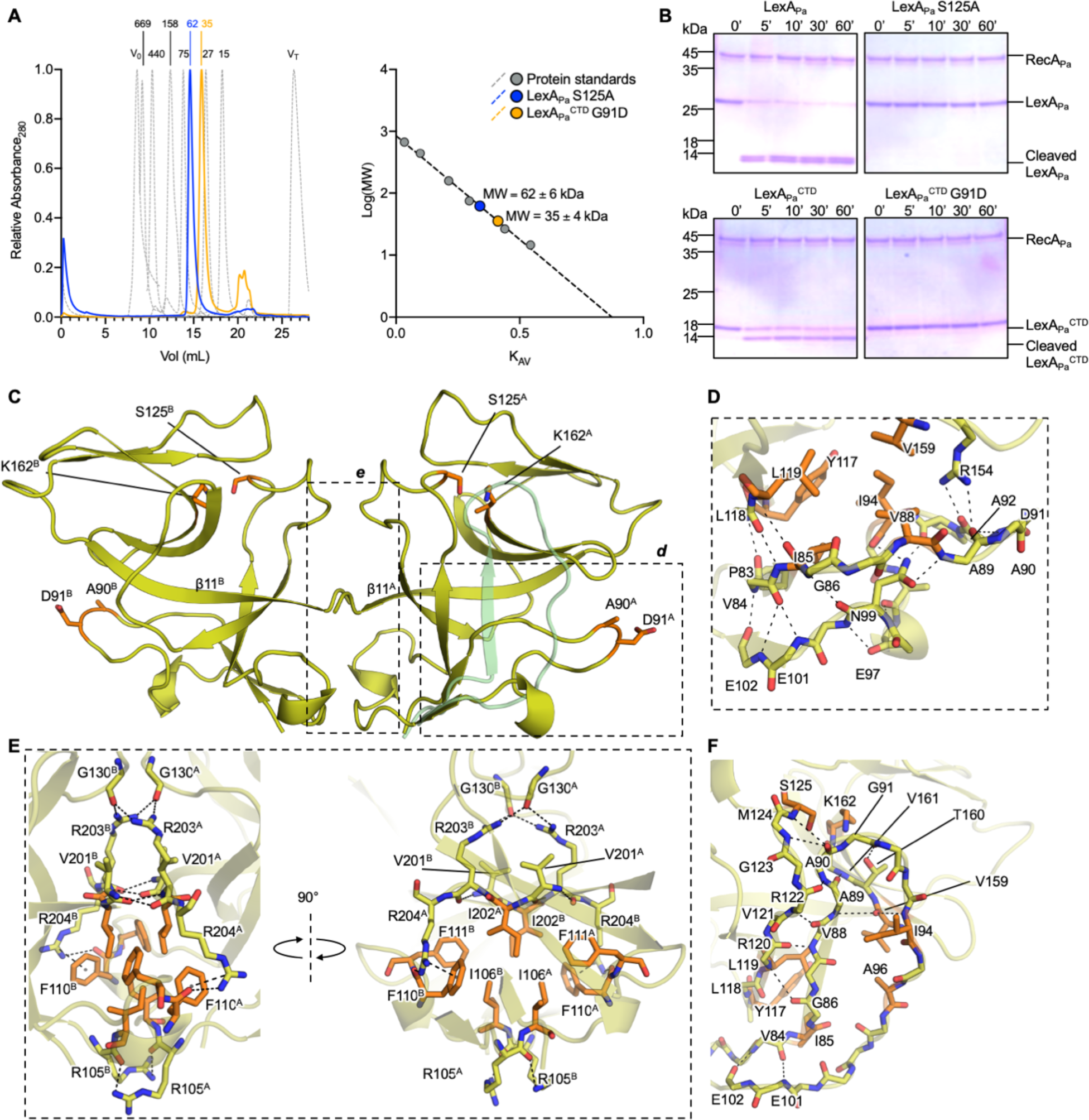
Structural analysis of LexA_Pa_^CTD^. (**A**) Analytical size exclusion chromatography of LexA_Pa_S125A (blue) and LexA_Pa_^CTD^G91D (yellow; chromatograms on the left and standard curve interpolation on the right). (**B**) SDS-PAGE-based RecA_Pa_*-induced autoproteolysis assay of 4 LexA_Pa_ variants: full-length LexA_Pa_, either wt or S125A inactive mutant, and LexA_Pa_^CTD^, either wt or G91D uncleavable mutant. (**C**) Overall view of the LexA_Pa_^CTD^G91D dimer (chains A and B), as revealed by X-ray crystallography. The catalytic dyad (S125/K162) and the mutated self-cleavage site (A90-D91) of each monomer are shown as orange sticks. Boxed regions are zoomed in panels D and E. Superposed (transparent green cartoon) is the closed conformation of LexA_Pa_ cleavable loop found in LexA_Pa_S125A bound to RecA_Pa_*. (**D**) Detailed view of the cleavable loop (chain A) in the “open” (inactive) conformation. Hydrogen bonds engaging the residues of the loop are represented as dashed lines, while residues involved in a hydrophobic cluster are depicted as orange sticks. (**E**) Detailed views of the homodimerization surface of LexA_Pa_^CTD^. Dashed lines indicate H-bonds, salt bridges and cation-π interactions, while residues involved in a hydrophobic cluster are depicted as orange sticks. (**F**) LexA_Pa_ cleavable loop in the “closed” (active) conformation. Dashed lines indicate H-bonds stabilizing the loop in this state, while orange sticks correspond to the catalytic dyad and to the hydrophobic residues indicated in panel B. The movement of the loop brings the cleavage site inside the catalytic pocket and at the same time opens a hydrophobic cavity (Y117, L119, V159, I85) that hosts I94 in the open conformation.

The structure of LexA_Pa_^CTD^ G91D has been resolved by X-ray macromolecular crystallography at 1.70 Å resolution (PDB ID 8B0V; statistics in Supplementary Table 1). Two independent molecules of LexA_Pa_^CTD^ G91D define the asymmetric unit and are fully visible from residue Gly81 to Arg204, while the functional homodimer can be reconstructed by applying a crystallographic symmetry operator and is hereafter referred to as chains A and B (indicated as superscript; Fig 1 C).

Electron densities that could not be assigned either to protein or ordered solvent have been interpreted as two calcium cations, two MES molecules, and three ethylene glycol molecules, all components of the crystallization conditions and not involved in any functional contact with the protein. A few weak electron densities remain uninterpreted and may be due to traces of the Tb-Xo4 nucleating agent (Engilberge *et al*, 2017, 2019) used in the crystallization process.

The homodimerization of LexA_Pa_^CTD^ G91D is mainly driven by the antiparallel pairing of the C-terminal portion of the β11 strands (secondary structures are numbered in Supplementary Fig. 1 A) of the interacting protomers (Fig. 1 C and 1 E). More in detail, Val201^A^ backbone oxygen and nitrogen are hydrogen bonded to Arg203^B^ nitrogen and oxygen, respectively. Other hydrogen bonds are established between Arg203^A^-NH1 and Gly130^B^-O, Arg204^A^-NH1 and Phe110^B^-O, Arg105^A^-NH1 and Arg105^B^-O. The sidechain of Arg204^A^ forms a cation-*π* interaction with the aromatic ring of Phe110^B^. Since all these interactions are mutual, they appear twice at the interaction surface. The core of LexA_Pa_^CTD^ G91D homodimerization surface is further stabilized by a hydrophobic cluster involving Ile106, Phe110, Phe111, and Ile202 of each chain (Fig. 1 E).

The cleavage loop (residues 81-103) of both LexA_Pa_^CTD^ G91D chains is in the inactive “open” conformation, with the mutated cleavage site (Ala90-Asp91) distant from the catalytic pocket that hosts the dyad Ser125/Lys162. This conformation is similar to those assumed by previously crystallized LexA^CTD^ mutants from other bacterial species (e.g., PDB 1JHF, 3JSP, 3K2Z).

In the “open” conformation, the base of the cleavage loop (Pro83-Ile85) is structured as a β-strand and pairs parallel to the β-strand Leu118-Arg120 (three intrachain hydrogen bonds are established between the backbone atoms; Fig. 1 D). The other extremity of the loop (Ile100-Cys104) assumes a β-sheet structure as well, and pairs in an antiparallel fashion with the aforementioned strands. On the tip of the loop, the backbone oxygen atoms of Ala89 and Ala92 are hydrogen bonded to the η nitrogen atoms of Arg154, while Ile94 is buried among Val88, Ile85, Leu119, Tyr117, Val159, and Glu195, forming several hydrophobic interactions.

The conformation of LexA_Pa_^CTD^ G91D cleavable loop was compared to that of LexA_Pa_S125A, subsequently obtained by Cryo-EM in complex with RecA_Pa_* (see the section “Cryo-EM structure of the RecA_Pa_*-LexA_Pa_ complex”; Fig. 1 C). The latter is in the active “closed” conformation (analogous to the one observed in PDB 1JHE, 3JSO, 8GMS and 8TRG), with the cleavage site buried inside the catalytic pocket. In this form, the β-strand that precedes the cleavage site extends until Ala90, increasing the number of interactions with the other core β-strands. Notably, in this conformation, Ile94 becomes solvent-exposed, opening the hydrophobic pocket where it was hosted in the open state (Fig. 1 F).

The sequence of LexA_Pa_ shows a high degree of identity with that of *E. coli* LexA (LexA_Ec_; 64% identity; Supplementary Fig. 1 A). As a consequence, LexA_Pa_^CTD^ G91D has a highly conserved structural arrangement compared to LexA_Ec_ (Supplementary Fig. 1 B; RMSD of 0.98 Å between LexA_Pa_^CTD^ G91D and PDB 1JHF, calculated over 124 pairs of α-carbon atoms by Gesamt; Krissinel, 2012). However, LexA_Pa_ displays a shorter C-terminal tail and a longer linker region between its CTD and NTD than LexA_Ec_ and these differences should be considered in the rational design of potential inhibitors of LexA_Pa_.

### Cryo-EM structure of RecA_Pa_*

RecA_Pa_ was expressed in *E. coli* and purified by affinity chromatography. To assemble the active nucleoprotein complex, RecA_Pa_ was co-incubated with 72mer oligo(dT) ssDNA and the slowly hydrolysable adenine nucleotide ATPγS. The desired RecA_Pa_* oligomers were stabilized by chemical crosslinking and isolated by size exclusion chromatography before vitrification of samples for Cryo-EM analysis.

The Cryo-EM structure of RecA_Pa_* was obtained by helical reconstruction, at a global resolution of 4.2 Å (Fig. 2 A, Supplementary Fig. 2 and 4 A-B, and Supplementary Table 2; PDB ID 8S70, EMD-19761). The final RecA_Pa_* model is organized as a right-handed helix described by a twist of 59.2°, a rise of 15.4 Å, ∼six RecA_Pa_ protomers per turn (corresponding to a pitch of 92.5 Å), and an average diameter of ∼110 Å, similar to that reported for the *E. coli* homolog (preprint: Cory *et al*, 2023; Yang *et al*, 2020; Chen *et al*, 2008; Gao *et al*, 2023; Fig. 2 B-D and Supplementary Fig. 2). This arrangement shows the features of RecA/ssDNA filaments in the ATP-bound extended form (Bell & Kowalczykowski, 2016; VanLoock *et al*, 2003). The density allowed the assignment of residues 1-328, and the identification of the contact sites with ssDNA and ATPγS (Fig. 2 E-G).

**Fig. 2:**
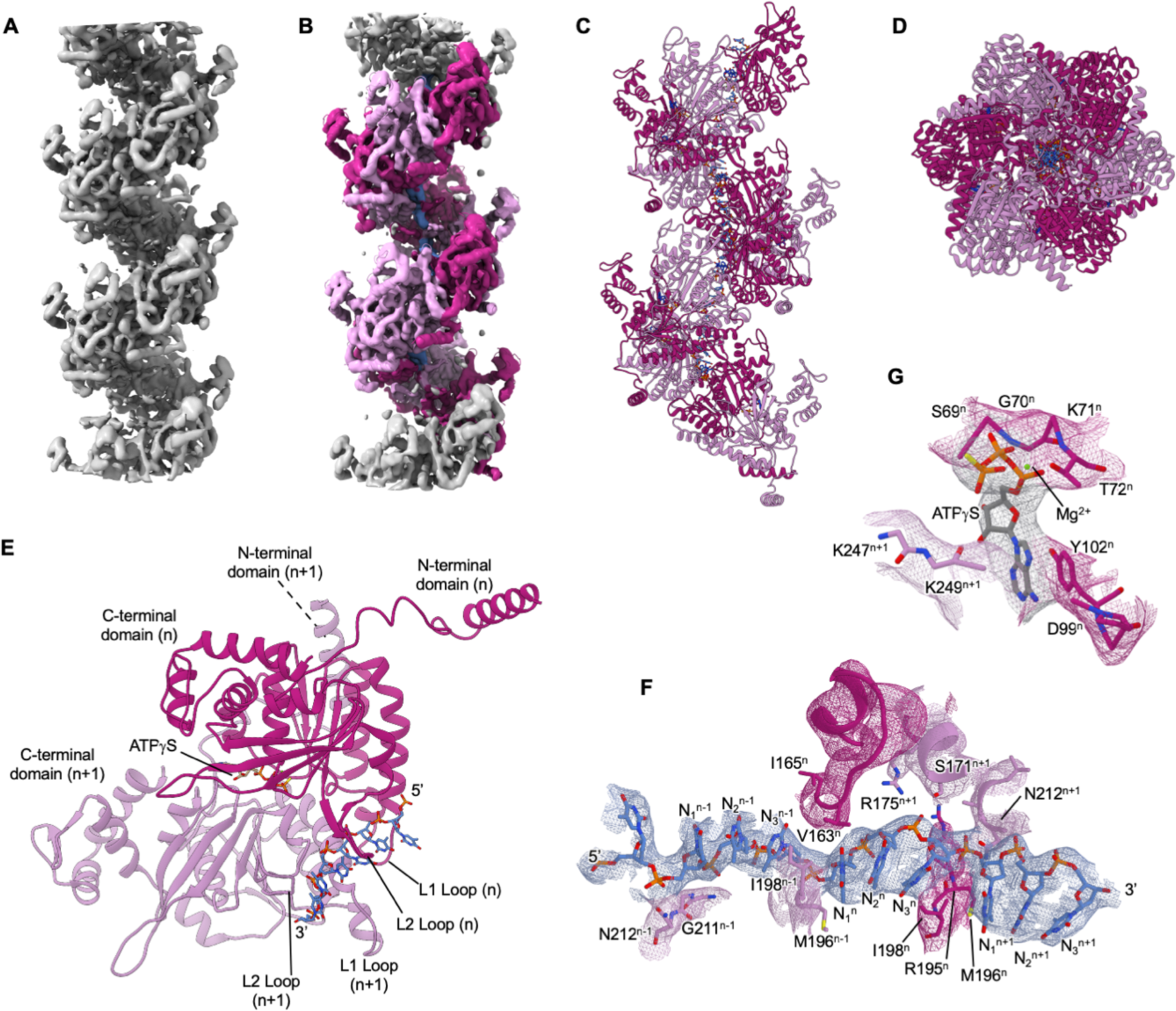
Cryo-EM structure of RecA_Pa_*. (**A**) RecA_Pa_* Cryo-EM density map. (**B**) Coloring of density regions corresponding to RecA_Pa_* protomers and (**C, D**) zoom on the atomic model (two perpendicular views). (**E**) Zoom on two adjacent RecA_Pa_* protomers assembled on ssDNA (RecA_Pa_^n^ and RecA_Pa_^n+1^, moving from 5’ to 3’ on ssDNA). Detailed views of the Cryo-EM map around ssDNA (**F**) and ATPγS (**G**), and RecA_Pa_ residues interacting with them.

Each RecA_Pa_ protomer interacts with the ssDNA filament by the central core domain (including seven α-helices and seven β-strands), from which the N- and C-terminal domains protrude. The N-terminal domain is constituted by a long α-helix and a flexible loop, while the C-terminal domain is mainly composed of three α-helices (α9-α11) and an intervening three-stranded β-sheet (β12-β14). The ssDNA, which lies close to the central axis and wraps around it, is contacted by RecA_Pa_ “ventral” L1 and L2 loops (residues 156-164 and 194-213; Fig. 2 E and Supplementary Fig. 1 C). The N-terminal helix of one RecA_Pa_ protomer (“*n+1*”) points toward the 5’ termini of ssDNA filament and docks on the “dorsal” part of the adjacent RecA_Pa_ monomer (“*n*”; Fig. 2 E), interacting with the α-helix 120-134 residues mainly by the formation of a cluster of hydrophobic residues (Leu114, Ile127, Leu131 and Val137 of RecA_Pa_^n^ and Leu9, Leu13, Ile16, Phe20 and Val25 of RecA_Pa_^n+1^). An average surface area of 2047 Å^2^ is buried on each RecA_Pa_ protomer at the interface with each of its neighboring ones, potentially establishing multiple van der Waals contacts and H-bonds.

One ATPγS molecule is coordinated at the interface between two RecA_Pa_ protomers (Fig. 2 G). Given the limited resolution of our maps, we can only speculate about the main interactions that this nucleotide might establish, by comparing the nucleotide binding pocket to previous structures of RecA_Ec_* oligomers obtained at higher resolution (PDB 7JY6 and 3CMW; Yang *et al*, 2020; Chen *et al*, 2008). ATPγS phosphate groups coordinate a Mg^2+^ cation, which in turn is kept in place by the side chain of Thr72 of RecA_Pa_^n^ (Fig. 2 G). The phosphate moieties are stabilized by hydrogen bonds with the backbone atoms of residues 68-73 of RecA_Pa_^n^, and by salt bridges with the side chains of Lys71^n^, Lys247^n+1^ and Lys249^n+1^. The adenine base might interact with acidic residues Asp99 of RecA_Pa_^n^, Asp249 and Glu250 of RecA_Pa_^n+1^ and can be further stabilized by interacting with Tyr102^n^.

When RecA_Pa_ is complexed with ssDNA, each RecA_Pa_ protomer spans mainly three nucleotides (5’-N_1_-N_2_-N_3_-3’; Fig. 2 F) but further contacts the phosphates of one nucleotide upstream (N_3_^-1^) and one nucleotide downstream (N_1_^+1^) of the primarily engaged triplet (Fig. 2 F). A physical torsion can be observed between nucleotides N_3_ and N_1_^+1^ (or, equivalently but in the opposite direction, between N_1_ and N_3_^-1^), with the side chain of Ile198^n^ inserting between their nucleobases. The phosphate group of N_1_ is at H-bond distance to RecA_Pa_^n^ Asn212 and Met196^n-1^, while the phosphate of N_2_ interacts with the backbone nitrogen atoms of Gly210^n^ and Gly211^n^. The negatively charged phosphate group of N_3_ could contact the side chains of Arg195^n^ and Arg175^n+1^, as well as Thr209^n^ and Ser171^n+1^. These interactions are encountered periodically along the RecA_Pa_* filament as they are established with the backbone of the DNA strand. Other local electrostatic or hydrophobic contacts with nucleobases depend on the nucleotide sequence.

RecA_Pa_ is highly similar to *E. coli* RecA (RecA_Ec_) in terms of both sequence (71% identity; Supplementary Fig. 1 C) and structure (RMSD of 1.06 Å between RecA_Pa_ chain F and PDB 7JY6 chain F, calculated over 320 pairs of α-carbon atoms by Gesamt; Supplementary Fig. 1 D), with the highest local differences affecting the C-terminal domain (residues 280-328), and the “ventral” loops (residues 159-165, 199-203 and 231-235).

### Cryo-EM structure of the RecA_Pa_*-LexA_Pa_ complex

To gain insights into the interaction between RecA_Pa_* (RecA_Pa_/ssDNA/ATPγS) and LexA_Pa_, the two interactors were co-incubated, chemically crosslinked, and the desired complexes were isolated by size exclusion chromatography for subsequent Cryo-EM studies. Since the interaction of LexA with RecA* triggers LexA autoproteolysis, to identify its docking site onto RecA_Pa_* but preventing hydrolysis occurrence, the LexA_Pa_S125A non-cleavable mutant was used to form the complex.

The structure of RecA_Pa_* in complex with LexA_Pa_S125A was determined by Cryo-EM at an overall resolution of 3.4 Å (Fig. 3 A, Supplementary Fig. 3 and 4 C-D, and Supplementary Table 2; PDB ID 8S7G, EMD-19771). Density for the LexA_Pa_ dimer is visible inside the helical groove of RecA_Pa_/ssDNA filament (Fig. 3 B). Interestingly, only the C-terminal domains (residues 81-204) of both LexA subunits were traceable in the maps, while the N-terminal DNA binding domains were largely undefined. A blurred extra density at low resolution (>7 Å) is observed protruding from the LexA_Pa_^CTD^ dimer. Although we cannot rule out the possibility that it derives from residual traces of map averaging, its position and size suggest it corresponds to the LexA_Pa_ NTD domain (Fig. 3 B and Supplementary Fig. 3). Its poorly defined nature is probably due to intrinsic flexibility, supporting the notion that the main binding determinants are located on the C-terminal domains, where the autocleavage reaction should occur.

**Fig. 3:**
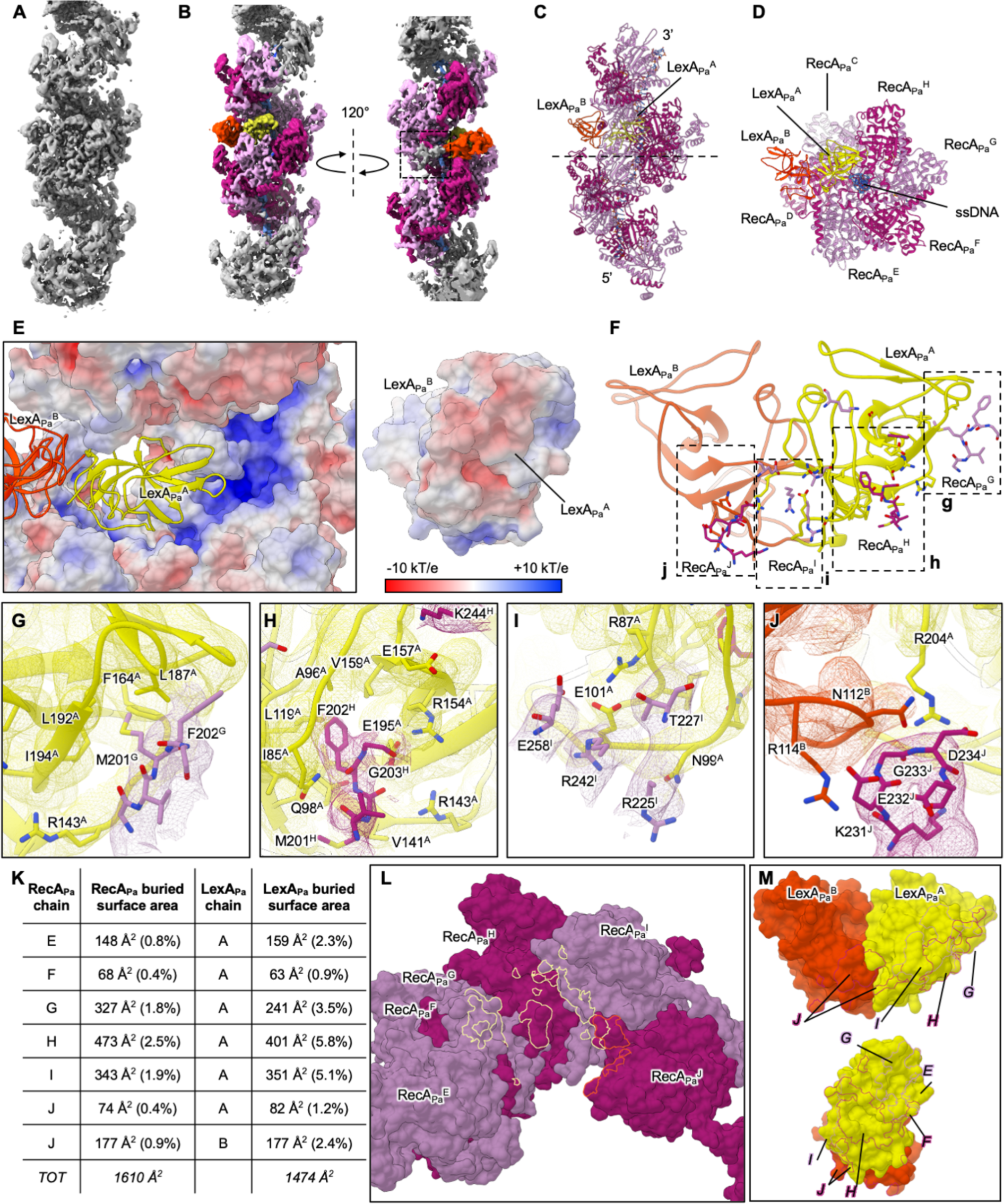
Cryo-EM structure of RecA_Pa_*-LexA_Pa_S125A. (**A**) Cryo-EM density map of the RecA_Pa_-LexA_Pa_S125A complex. (**B**) Coloring of density regions corresponding to RecA_Pa_* protomers (purple tones) and LexA_Pa_ CTD chains A (yellow) and B (orange). The boxed region represents a low-resolution density, which was not interpreted by the atomic model and that might be due to the LexA_Pa_ NTD. (**C**) Side and (**D**) front views of the RecA_Pa_*-LexA_Pa_S125A atomic model. The dashed line in panel **C** represents a virtual plane where the model was cut in panel **D** to allow LexA_Pa_ clear visualization. (**E**) Electrostatic surface potential of RecA_Pa_* and LexA_Pa_^CTD^, showing complementarity on the interacting surfaces. (**F**) LexA_Pa_^CTD^ dimer and the main binding determinants on four RecA_Pa_ protomers (chains G-J), zoomed in panels **G-J**. (**K**) Details of the interfaces buried between LexA_Pa_ and different RecA_Pa_* protomers. The corresponding interacting surfaces are represented in panels **L** (on RecA_Pa_* surface) and **M** (on LexA_Pa_ surface, front and side views). Contour lines are colored as the interacting chain.

Our data showed that full-length LexA_Pa_ non-stoichiometrically occupies the RecA_Pa_* helical groove and hence does not follow its helical symmetry (Fig. 3 B-D). Indeed, the RecA_Pa_*-LexA_Pa_ complex could only be resolved by local refinement after dropping the helical symmetry assumption. The LexA_Pa_^CTD^ subunit deeply buried inside RecA_Pa_* groove (chain A or LexA_Pa_^A^) is well defined in the maps and directly interacts with three consecutive protomers of RecA_Pa_ (chains G, H and I, assembled 3’-5’ on ssDNA), contacting the L2 loops (residues 197-207) of two of them (G and H) and the core β-strands of the third (Fig. 3 E-M). Conversely, chain B (LexA_Pa_^B^, which is slightly less well-defined in the map) keeps the LexA dimeric arrangement, but remains more peripheral, most likely establishing a few contacts only with RecA_Pa_* chain J. The cleavable loop of LexA_Pa_ chain A assumes the closed conformation, producing a hydrophobic cavity (defined by the residues Ile85, Ala96, Tyr117, Leu119, Leu137, Val139, Val152, Val159, and Glu195) that is explored by Phe202 of RecA_Pa_ chain H (Fig. 3 H). This complex architecture suggests that the L2 loop of RecA_Pa_^H^ (residues 202-204) is kept in place by a network of polar interactions established at the interface between RecA_Pa_^H^ Met201, Phe202 and Gly203 and side chains of LexA_Pa_^A^ Gln98, Arg143, Arg154 and Glu195. LexA_Pa_^A^ might establish additional contacts with the L2 loop of the upstream RecA_Pa_ protomer in the helical assembly (RecA_Pa_^G^): in this case Met201 protrudes into a nearby hydrophobic pocket of LexA_Pa_^A^, defined by Leu187, Leu192, Phe164 and Ile194 (Fig. 3 G). LexA_Pa_ chain B maintains its cleavable loop in the open state as in the X-ray structure of LexA_Pa_^CTD^ G91D described above, with the hydrophobic pocket made inaccessible by LexA_Pa_ Ile94.

Analyzing the surface electrostatic potential of each member of the complex, the groove of RecA_Pa_* oligomer (positively charged) and the interacting flank of LexA_Pa_^A^ in the cleavable conformation (negatively charged) display a wide and remarkable complementarity (Fig. 3 E). Even though the map resolution does not allow to clearly define the position of their side chains, electrostatic interactions are likely established between RecA_Pa_^H^ Lys244, RecA_Pa_^I^ Lys231, RecA_Pa_^I^ Arg242, RecA_Pa_^I^ Glu258 and RecA_Pa_^J^ Asp234 and LexA_Pa_^A^ Glu157, Glu158, Glu101, Arg87 and Arg204 (Fig. 3 I-J).

### Fluorescence polarization-based analysis of RecA_Pa_ affinity for its ligands

To characterize the binding affinity of RecA_Pa_ for its ligands (ssDNA and ATPγS), a fluorescence polarization (FP)-based assay was set up using a Fluorescein amidite (FAM)-labeled 32mer ssDNA filament and leveraging on the FP increase observed upon RecA_Pa_ oligomerization on it (Lee *et al*, 2007).

A first experiment (Fig. 4 A) was carried out by titrating ssDNA with RecA_Pa_ in the presence of a large excess of ATPγS. The apparent affinity of RecA_Pa_ for FAM-32mer ssDNA, resulting from data fitting with the Hill equation, lies in the nanomolar range (*K_D_*^App^ = 82 ± 34 nM) and the Hill coefficient suggests binding cooperativity (*h* = 1.9 ± 0.3), as expected for RecA oligomerization on DNA (Cory *et al*, 2022).

**Fig. 4:**
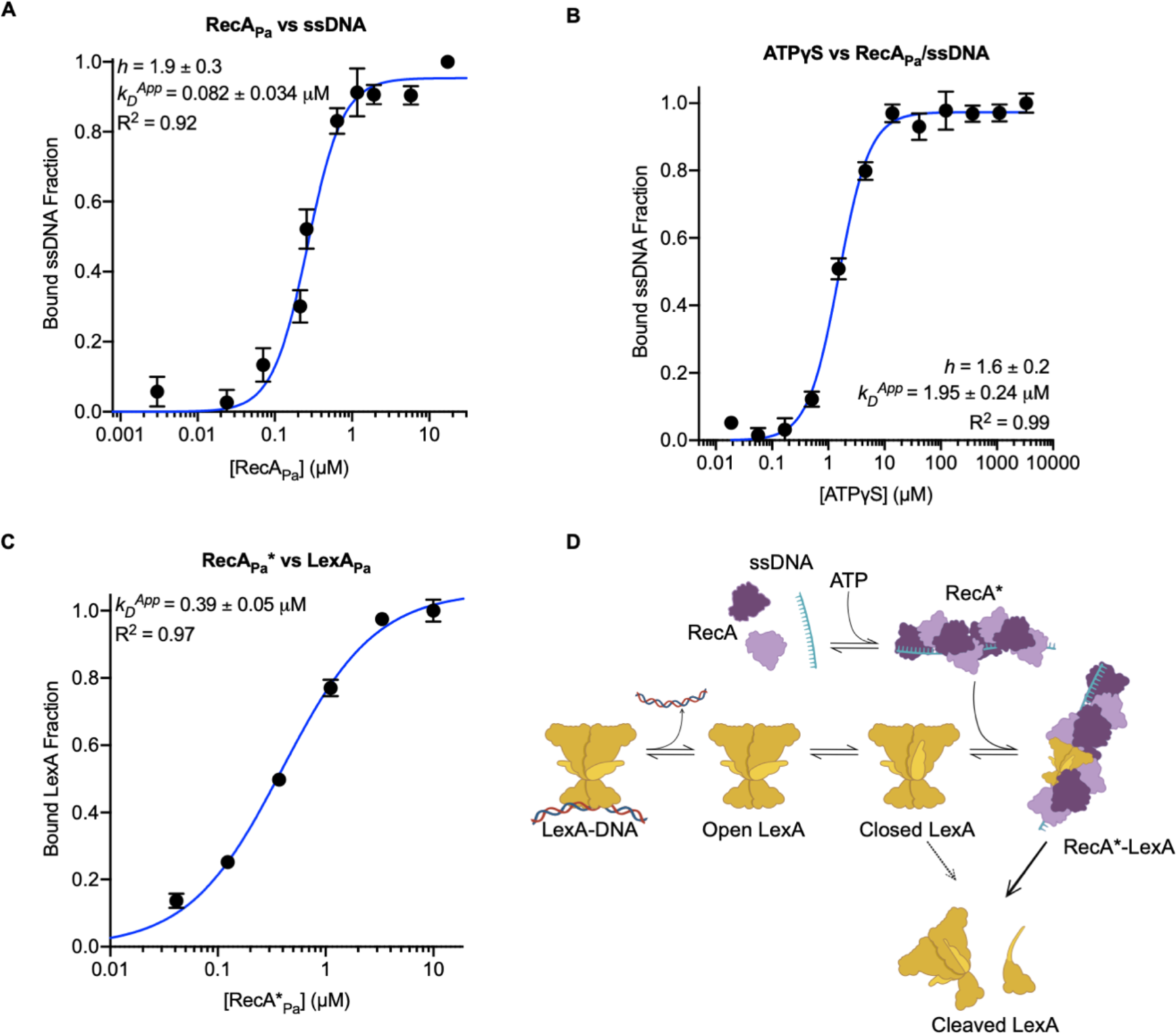
Analysis of RecA_Pa_ interactions with its natural ligands (ATP*γ*S, ssDNA and LexA_Pa_). FP-based titrations of (**A**) FAM-32mer ssDNA with RecA_Pa_ (ATP*γ*S in molar excess), (**B**) RecA_Pa_/FAM-32mer ssDNA with ATP*γ*S and (**C**) FlAsH-LexA_Pa_^CTD^S125A with activated RecA_Pa_ (RecA_Pa_*, RecA_Pa_/ssDNA/ATP*γ*S). Points represent the average of three replicates while error bars represent standard errors. (**D**) Overview of the model proposed for the molecular process promoted by RecA_Pa_*, that leads to the autocleavage of LexA_Pa_. LexA_Pa_ can bind RecA_Pa_* if it is free from DNA and with the cleavable loop in the closed conformation. The binding to RecA_Pa_* allows the self-cleavage of LexA_Pa_, that otherwise is mainly prevented.

A second experiment (Fig. 4 B) was performed by keeping the concentration of FAM-32mer ssDNA and RecA_Pa_ constant while increasing the concentration of ATPγS in the different samples. The obtained apparent *K_D_* of RecA_Pa_ for ATPγS in the presence of FAM-32mer ssDNA (*K*_D_^App^ = 2.0 ± 0.2 µM; *h* > 1.5) is about 5 times higher than the previously determined *K_D_* of *E. coli* RecA for the same nucleotide (K_D_ < 0.4 µM; Kowalczykowski, 1986). Such differences might arise from structural peculiarities of *E. coli* and *P. aeruginosa* RecA ATP-binding sites or from limitations of the different experimental methods used. In particular, the biophysical assay reported here indirectly measures the affinity of RecA_Pa_ for ATPγS, since it relies on RecA_Pa_ oligomerization on the fluorescent reporter FAM-32mer ssDNA and on the influence that the nucleotide cofactor has on this interaction.

A different FP-based assay was employed to estimate the affinity of RecA_Pa_* for LexA_Pa_ (Fig. 4 C), titrating a fluorescently labeled and uncleavable variant of LexA_Pa_ C-terminal domain (FlAsH-LexA_Pa_^CTD^ S125A) with increasing concentrations of pre-activated RecA_Pa_*. The latter was oligomerized on a 18mer ssDNA, to provide a single but fully functional LexA_Pa_ binding site. A high-nanomolar dissociation constant has been obtained by fitting the data using a bimolecular binding model (*K*_D_^App^ = 390 ± 50 nM).

## Discussion

In this work, we solved the structure of the main regulative players of the SOS response (RecA and LexA) in *P. aeruginosa,* a relevant human pathogen whose DNA-damage response still requires thorough understanding.

The structures of LexA_Pa_ C-terminal autoproteolytic domain (Fig. 1), activated RecA_Pa_ (i.e., complexed with ssDNA and ATPγS, referred to as RecA_Pa_*; Fig. 2), and their complex (RecA_Pa_*-LexA_Pa_S125A; Fig. 3) were obtained by X-ray crystallography and Cryo-electron microscopy, respectively.

While writing our manuscript, we came across a preprint publication describing the complex of *E. coli* RecA and full-length LexA (preprint: Cory *et al*, 2023). Besides supporting our main results, the structure detailed by Cory and colleagues, together with previous structural studies (Gao *et al*, 2023), offered the opportunity to highlight peculiarities of the SOS components and activation complex here disclosed.

RecA_Pa_ has been structurally investigated in complex with ssDNA and ATPγS, showing an extended helical assembly, which is kept unaltered upon LexA_Pa_ binding.

RecA_Pa_ sequence is highly homologous to RecA_Ec_, with the highest differences affecting the very C-terminal tail (residues 330-346 in *P. aeruginosa*; Supplementary Fig. 1 C). In both species this region has a high percentage of acidic residues and is likely very flexible, thus it is not visible in previous (e.g.: PDB 7JY6 and 3CMW; Yang *et al*, 2020; Chen *et al*, 2008) and in our structures. Superposition of RecA_Pa_* structure to RecA_Ec_* (PDB 7JY6; Yang *et al*, 2020) revealed a very high global and local structural similarity, with conservation of the main residues defining the binding sites for ATPγS and ssDNA.

The repressor LexA_Pa_, whose C-terminal autoproteolytic domain has been resolved at 1.70 Å resolution, displays a dimeric assembly with a fully resolved cleavable loop in the open conformation in the crystal packing. When compared to the LexA_Ec_ homolog, the full-length LexA_Pa_ displays a shorter C-terminal tail and a longer linker region between its CTD and NTD. Both these regions might contribute to notable binding sites of regulators or putative inhibitors, given their proximity to the cleavage loop. For instance, these areas of LexA_Ec_ are involved in the binding of both phage GIL01 gp7 LexA-modulating protein (Caveney *et al*, 2019) and recently developed anti-LexA nanobodies (Maso *et al*, 2022).

The SOS complex of *P. aeruginosa,* here described for the first time, reveals a unique architecture: the full-length LexA_Pa_S125A decorates RecA_Pa_* non-stoichiometrically. This was clearly confirmed by our data processing, as all the attempts made to reconstruct the complex using helical refinement, imposing RecA_Pa_* helical symmetry, failed. On the other hand, using single particle reconstruction and homogenous refinement, we obtained a clear and well-defined density (Fig. 3 B), corresponding to a dimer of LexA_Pa_S125A C-terminal domains into the groove of a six-member turn of RecA_Pa_* (Fig. 3 C-D). This result agrees with the *E. coli* complex recently described by Cory and coworkers (preprint: Cory *et al*, 2023), while it disagrees with the previous RecA_Ec_*-LexA_Ec_^CTD^ complex structure (PDB 8GMS; Gao *et al*, 2023), where LexA autoproteolytic CTD was fully decorating the RecA_Ec_* filament and followed its helical symmetry. The symmetrical architecture observed by Gao and colleagues is likely due to the absence of LexA_Ec_ NTD domains, which cannot exert any steric hindrance on adjacent LexA binding sites.

Full-length LexA_Pa_ binding mainly entails the engagement of three consecutive RecA_Pa_ protomers (chains G, H and I; Fig. 3 F-I), as shown by the extension of the buried surface areas: 993 Å^2^ are buried on LexA_Pa_^A^ (14.4 % of its total surface) at the interface with these three chains (Fig. 3 K-M). Among these three, the central one (chain H in our complex) contributes most and protrudes with Phe202 (located on the L2 loop) in a hydrophobic pocket that is formed only upon closure of LexA_Pa_ cleavable loop towards its catalytic crevice (Fig. 1 F and Fig. 3 H). Here, several polar and non-polar interactions can stabilize the two binding partners. The upstream RecA_Pa_ protomer (chain G; towards the 3’ terminus of ssDNA) contacts the same LexA_Pa_ chain by hydrophobic/van der Waals interactions, while the downstream RecA_Pa_ protomer (chain I; towards the 5’ terminus of ssDNA) could define multiple polar contacts with LexA_Pa_^A^ (Fig. 3 G-I). Last, a fourth RecA_Pa_ protomer (chain J) is placed at a distance compatible with further electrostatic interactions with both chains of the LexA_Pa_ dimer (Fig. 3 J). However, the contribution of chain J to the binding of LexA_Pa_ is likely very limited, as noticed by Cory and colleagues for *E. coli* (preprint: Cory *et al*, 2023).

The protein surface and key determinants of the RecA*-LexA interaction are highly conserved between *E. coli* and *P. aeruginosa*. A peculiar difference consists in the conformation of the LexA repressor NTD domain. Indeed, in both complexes it partially occupies the groove of RecA* but it results poorly defined and more peripheral in the *P. aeruginosa* structure. Such differences are most likely due to a roughly twice longer linker connecting the NTD and CTD domains of LexA_Pa_ (eleven versus five amino acids of the *E. coli* homologue). Such a long spacer introduces higher flexibility between the two domains of LexA_Pa_ and it might prevent the formation of stable interactions by the NTD domain with RecA_Pa_* oligomers.

The structures here presented unravel that the main determinants of the activation process reside in the CTD domain, supporting the significance of the RecA_Pa_*-LexA_Pa_^CTD^ binding measurements performed *in vitro* on recombinant purified species (Fig. 4). Dissociation constants of RecA_Pa_ to ssDNA (to form RecA_Pa_*) and RecA_Pa_* to LexA_Pa_^CTD^, evaluated by dedicated FP-based assays, fall in the mid (*K*_D_^App^= 82 ± 34 nM) and high nanomolar range (*K*_D_^App^= 390 ± 50 nM), respectively. Despite it might be affected by the oligonucleotide length used in the assay, the affinity of RecA_Pa_ towards ssDNA is in the expected range. The binding constant between the components of the activation complex results in a remarkable agreement with the previously determined one for a full-length *E. coli* LexA S119A with its cognate RecA* (360 nM; Cory *et al*, 2022).

Our experimental data strongly supports the most accepted model proposed for the activation of the SOS response (Fig. 4 D). In the absence of *‘SOS’ stimuli*, the equilibrium between the closed and the open conformations of LexA cleavage loop largely favors the uncleavable one, leaving LexA intact and capable of repressing the SOS genes. After exposure to stressors, the resulting DNA damage promotes RecA* nucleoprotein filaments assembly, providing a molecular surface able to selectively bind LexA in the closed conformation and free from dsDNA (as SOS box DNA is known to hamper RecA* binding; Butala *et al*, 2011). This binding event alters the equilibrium between LexA conformations in favor of the cleavable one, while co-catalyzing the LexA autocleavage.

This notion finds a clear support in the structural analysis of the complex, where only the closed state of LexA fits the binding region of RecA* oligomers, and the cleavable loop is engaged in extensive interactions with recombinase protomers by residues distributed both upstream and downstream the scissile peptide bond.

Since LexA cleavable loop contributes to define the hydrophobic pocket that hosts RecA key phenylalanine, upon LexA autoproteolysis the binding site for RecA* is divided among the cleavage products. It is likely that this allows their dissociation from RecA*. This model agrees with previous observations that LexA^CTD^ affinity for RecA* remains comparable to that of full-length LexA, provided that the N-terminal truncation leaves intact the initial structured region of the CTD (starting at residue G75 in *E. coli,* G81 in *P. aeruginosa*; Hostetler *et al*, 2020).

A deep understanding of the SOS response at the molecular level is of great significance for both general and medical microbiology. Indeed, this stress response pathway to DNA damage is widely recognized as one of the main drivers of the evolution of antibiotic resistance and a master regulator of several disease-related phenomena. On the other hand, recent works have pinpointed significant inter-species differences in this conserved and long-studied pathway, underlining that it still has hidden aspects, especially in non-model organisms.

The structures of the essential SOS components and their activation complex in the *P. aeruginosa* pathogen, as presented here, along with the recent models revealed for the *E. coli* bacterial model, have successfully addressed a gap that persisted for over three decades in basic research. These findings have uncovered pivotal elements, crucial for designing innovative strategies to combat bacterial pathogens, focusing on anti-evolutionary and antivirulence approaches.

## Author contributions

Filippo Vascon: Conceptualization; Investigation; Formal analysis; Data curation; Methodology; Writing - original draft, review and editing. Sofia De Felice: Conceptualization; Investigation; Formal analysis; Data curation; Methodology; Writing – original draft, review and editing. Matteo Gasparotto: Conceptualization; Investigation; Formal analysis; Data curation; Writing - original draft. Stefan Huber: Methodology; Formal analysis; Data curation. Monica Chinellato: Investigation; Formal analysis; Methodology. Claudio Catalano: Methodology; Formal analysis; Data curation. Alessandro Grinzato: Methodology; Data curation. Francesco Filippini: Supervision; Data curation; Writing – review and editing. Lorenzo Maso: Investigation; Methodology; Formal analysis. Arjen Jakobi: Methodology; Formal analysis; Data curation. Laura Cendron: Conceptualization; Investigation; Writing – original draft; Methodology; Writing – review and editing; Project administration; Resources; Supervision; Validation.

## Acknowledgments

The authors would like to thank the staff of beamline ID30b of the European Synchrotron Radiation Facility (ESRF, Grenoble, France), Gregory Effantin and the staff at CM01 Microscopy beamline for technical assistance during data collections. We thank Giancarlo Tria for technical assistance at the FloCen facility in Florence. We acknowledge the support of the Italian Ministry of Education and Instruct-ERIC (PID6440, VID12221), part of the European Strategy Forum on Research Infrastructures (ESFRI). We thank EMBO Society for supporting Sofia De Felice with EMBO Scientific Exchange Grant Initiative (Grant Number 10252). We thank FEMS for supporting Filippo Vascon with a FEMS Research and Training Grant (FEMS-GO-2021-002). We thank Fondazione Cassa di Risparmio di Padova e Rovigo (Ca.Ri.Pa.Ro.) for supporting Filippo Vascon and Matteo Gasparotto with PhD scholarships.

## Conflict of interest statement

The authors declare that the research was conducted in the absence of any commercial or financial relationships that could define a potential conflict of interest.

## Materials and methods

### Molecular cloning and site-directed mutagenesis

The genomic DNA of *P. aeruginosa* ATCC 27853 was purified from an overnight liquid culture using the GenElute Bacterial Genomic DNA Kit (Merck) according to manufacturer’s instructions. The coding sequences of *P. aeruginosa* RecA (RecA_Pa_) and LexA C-terminal domain (LexA_Pa_^CTD^) were PCR-amplified from *P. aeruginosa* ATCC 27853 gDNA using primers RecA_Pa_pColi.For/Rev and LexA_CTD_Pa_pColi.For/Rev, respectively (Supplementary Table 3) and cloned in the pColiExpressI plasmid vector (Canvax) by ligation-independent cloning following manufacturers’ instructions. The obtained plasmids were named pColiXP-RecA_Pa_ and pColiXP-LexA_Pa_^CTD^. The coding sequence of *P. aeruginosa* full-length LexA (LexA_Pa_) and TetraCys-tagged LexA_Pa_^CTD^ were amplified from the genomic DNA using primers LexA_Pa.For/Rev and LexA_Pa_CTD_4Cys.For/Rev (Supplementary Table 3) and cloned in pETite C-His Kan vector and pETite N-His SUMO Kan Vector (Lucigen), respectively, following manufacturer’s instructions. The obtained plasmid vectors will be referred to as pETite-LexAPa and pETite-SUMO-4Cys-LexA_Pa_^CTD^.

The three plasmids encoding LexA_Pa_ variants were used as templates to introduce inactivating mutations either altering LexA_Pa_ cleavable loop (G91D) or its catalytic site (S125A), using the QuikChange Site-Directed Mutagenesis kit (Agilent Technologies) and mutagenic primers listed in Supplementary Table 3.

All generated plasmids were verified by Sanger sequencing.

### Recombinant protein expression and purification

#### RecA_Pa_

N-terminal His-tagged *P. aeruginosa* Recombinase A (RecA_Pa_) was expressed in *E. coli* BL21(DE3) cells, transformed with pColiXP-RecA_Pa_ and grown in LB broth supplemented with 100 μg/mL ampicillin. Protein overexpression was induced by adding 1 mM isopropyl-β-D-thiogalactoside (IPTG) to the bacterial culture in the late exponential growth phase (OD_600_ 0.6-0.8) and was carried out overnight at room temperature under vigorous shaking (180 rpm). Cells were harvested by centrifugation and resuspended in buffer R_A (10 mM HEPES, 300 mM NaCl, 10% v/v Glycerol, 20 mM Imidazole, pH 8.0) supplemented with 1X Protein Inhibitors Cocktail (SERVA) and a tip of spatula of DNAse I (Sigma Aldrich). Bacterial cells lysis was performed by sonication. Cell debris were removed by centrifugation and the lysate soluble fraction was loaded on a 5 mL HisTrap Excel IMAC column (Cytiva). His-tagged RecA_Pa_ was eluted after extensive buffer R_A washes, by linearly raising the imidazole concentration in the eluent from 50 mM to 500 mM in 3 column volumes. IMAC fractions showing RecA_Pa_ as the main protein component in SDS-PAGE analysis were pooled together, concentrated using a Vivaspin Turbo Ultrafiltration unit (10 kDa MWCO; Sartorius) and buffer-exchanged in 10 mM HEPES, 300 mM NaCl, 10% Glycerol, 1 mM MgCl_2_, 1 mM dithiothreitol (DTT), pH 7.0, by a HiTrap Desalting column (Cytiva) before storage at −80 °C for future usage in *in vitro* assays. Since the N-terminal 6xHisTag did not interfere with RecA_Pa_ assembly on ssDNA and with RecA_Pa_*-mediated LexA_Pa_ self-cleavage, it was not removed after protein purification.

#### LexA_Pa_ variants

N-terminal His-tagged LexA_Pa_, either wild-type or S125A catalytically-inactive mutant (LexA_Pa_S125A), and C-terminal His-tagged LexA_Pa_ C-terminal domain, either wild-type or G91D uncleavable mutant (LexA_Pa_^CTD^G91D), were expressed in *E. coli* BL21(DE3) cells, transformed with pETite-LexA_Pa_ (S125A) and pColiXP-LexA_Pa_^CTD^ (G91D), respectively. Cells were grown in LB broth supplemented with 50 μg/mL kanamycin or 100 μg/mL ampicillin, respectively. Protein overexpression was induced by adding 1 mM isopropyl-β-D-thiogalactoside (IPTG) to bacterial cultures in the late exponential growth phase (OD_600_ 0.6-0.8) and was carried out overnight at room temperature under vigorous shaking (180 rpm). Cells were harvested by centrifugation and resuspended in buffer L_A (20 mM Tris-HCl, 150 mM NaCl, 10 % v/v Glycerol, pH 7.5) supplemented with 20 mM Imidazole, 1X Protein Inhibitors Cocktail (SERVA), 500 U of benzonase nuclease (Merck) and 1.5 mM MgCl_2_. Bacterial cells were lysed by sonication and the crude lysate was incubated 30 minutes at 4 °C to allow benzonase-mediated DNA digestion. The supernatant was cleared by centrifugation and loaded on a 1 mL HisTrap Excel IMAC column (Cytiva). After thoroughly washing the resin with buffer L_A and with 20 mM imidazole in buffer L_A, His-tagged LexA_Pa_ variants were eluted by linearly raising the imidazole concentration in the eluent from 20 mM to 500 mM in 10 column volumes. IMAC fractions showing LexA_Pa_S125A as the main protein component by SDS-PAGE analysis were pooled together, concentrated using a Vivaspin Turbo Ultrafiltration unit (5 kDa MWCO; Sartorius) and buffer-exchanged to buffer L_A by a HiPrep 26/10 desalting column (Cytiva) before storage at −80 °C. IMAC fractions containing mostly pure 6His-LexA_Pa_^CTD^ G91D, as evidenced by SDS-PAGE analysis, were pooled together, concentrated and further purified by size-exclusion chromatography on a HiLoad Superdex 75 26/60 PG column (Cytiva) equilibrated in 20 mM tris-HCl pH 7.6, 150 mM NaCl, 5% v/v glycerol. The affinity tag was cleaved from 6His-LexA_Pa_^CTD^ G91D by incubating the purified protein overnight at 4 °C with recombinant TEV protease (LexA:TEV ratio of 20:1, w/w), 0.4 mM DTT, 0.15 mM EDTA and 0.01% v/v NP-40. The following day, the mixture was diluted twice with buffer L_A to reduce DTT and EDTA concentration and then loaded on a 1 mL HisTrap Excel IMAC column, recovering the flowthrough that contains LexA_Pa_^CTD^ G91D without 6xHisTag. This sample was then buffer exchanged to 20 mM tris-HCl pH 7.6, 150 mM NaCl, 5% v/v glycerol and concentrated to 11.5 mg/mL before storage at −80 °C for protein crystallization.

#### FlAsH-LexA ^CTD^S125A

N-terminal His-SUMO-4Cys-tagged *P. aeruginosa* LexA C-terminal domain S125A uncleavable mutant (6His-SUMO-4Cys-LexA_Pa_^CTD^S125A) was expressed in *E. coli* BL21(DE3) cells, transformed with pETite-SUMO-LexA_Pa_^CTD^S125A and grown in LB broth supplemented with 50 μg/ml kanamycin. Protein overexpression was induced by adding 1 mM isopropyl-β-D-thiogalactoside (IPTG) to the bacterial culture in the late exponential growth phase (OD_600_ 0.6-0.8) and was carried out overnight at room temperature under vigorous shaking (180 rpm). Cells were harvested by centrifugation and resuspended in buffer FL_A (20 mM Tris-HCl, 150 mM NaCl, 10 % v/v Glycerol, 0.1 mM DTT, pH 7.5) supplemented with 20 mM Imidazole and 1X Protein Inhibitors Cocktail (SERVA). Bacterial cells lysis was performed by sonication. Cell debris were removed by centrifugation and the lysate soluble fraction was loaded on a 1 mL HisTrap Excel IMAC column (Cytiva). After extensively washing the column with buffer FL_A and with 20 mM imidazole in buffer FL_A, 6His-SUMO-4Cys-LexA_Pa_^CTD^S125A was eluted by linearly raising the imidazole concentration in the eluent from 20 mM to 500 mM in 10 column volumes. IMAC fractions showing 6His-SUMO-4Cys-LexA_Pa_^CTD^S125A as the main protein component by SDS-PAGE analysis were pooled together, diluted three times in buffer FL_A and supplemented by 1 mM DTT, 1 mM EDTA, 0.1% v/v NP-40, and an excess of Expresso Sumo Protease (Lucigen). Following a 2-hours incubation at room temperature with gentle shaking, 100 μM FlAsH-EDT_2_ was added to the reaction mix and the incubation was prolonged overnight at 4 °C in the dark. The mixture was then concentrated using a Vivaspin Turbo Ultrafiltration unit (5 kDa MWCO; Sartorius) and buffer-exchanged to 20 mM tris-HCl, 150 mM NaCl, 10% v/v Glycerol, pH 7.5 by a PD-10 desalting column (Cytiva). To remove 6His-SUMO fragments and uncleaved protein constructs from the final sample, the mixture was passed through a 1 mL HisTrap Excel IMAC column (Cytiva) and the flowthrough was recovered. FlAsH-LexA_Pa_^CTD^S125A was stored at −80 °C for future usage in *in vitro* assays.

### SDS-PAGE-based RecA_Pa_*-mediated LexA_Pa_ autoproteolysis assay

RecA_Pa_ was co-incubated 1h at 37 °C with SKBT25-18mer ssDNA* ([RecA_Pa_]:[18mer ssDNA]=3.5:1) and a molar excess (1 mM) of ATPγS. To test the RecA_Pa_*-induced autoproteolytic activity of purified LexA_Pa_ variants, 1 μM of each variant was incubated with 1 μM RecA_Pa_* at 37 °C. 30 mM HEPES, 150 mM NaCl, pH 7.1 was used as the reaction buffer. The reaction was stopped at different time points by adding Laemmli sample buffer and incubating the samples 5 min at 95 °C before loading them on Bis-Tris-SDS 4-20% polyacrylamide gels (SurePAGE, GenScript).

### LexA ^CTD^G91D crystallization

11.5 mg/mL LexA_Pa_^CTD^ G91D underwent large-scale crystallization trials by the sitting-drop isothermal vapor diffusion method. 0.4 µL drops were produced mixing an equal volume of protein and precipitant solutions (PACT, LMB and JCSG-plus crystallization kits; Molecular Dimensions) by an Oryx8 dispensing robot (Douglas Instruments) and incubated at 293 K. The best crystals grew in buffers 1-23, 2-9 and 2-11 of the PACT premier crystallization trial kit and were further optimized by the addition of 5 mM Tb-Xo4 crystallophore (Polyvalan; Engilberge *et al*, 2017, 2019) to the protein solution as nucleating agent. Crystals were cryo-protected by adding 30% v/v PEG 400 to the mother liquor before freezing in liquid nitrogen for shipment to the synchrotron facility.

### X-ray structure determination

X-ray diffraction experiments of protein crystals were performed at the ID30B beamline of the European Synchrotron Radiation Facility (ESRF, Grenoble, France). Best LexA_Pa_^CTD^ G91D diffracting crystals were obtained in PACT 1-23 precipitant buffer (0.2 M CaCl_2_•2H_2_O, 0.1 M MES pH 6.0, 20% w/v PEG 6000). Collected data were analyzed by the available automated processing pipelines for space group determination and reflections indexing. Data reduction was performed by Aimless via the CCP4i2 interface (Evans, 2011). Molecular replacement was carried out by Molrep (Vagin & Teplyakov, 2010), using a homology model of LexA_Pa_^CTD^G91D generated by SwissModel (Waterhouse *et al*, 2018) using PDB 1JHF as a template. The protein model was adjusted by manual and automated structure refinement, using Coot (Emsley *et al*, 2010) and Refmac5 (Murshudov *et al*, 2011) via the CCP4i2 interface (Winn *et al*, 2011). The LexA_Pa_^CTD^G91D dimer was reconstructed in Pymol v2.3.5 applying the crystallographic symmetry operator.

### Isolation of multi-protein complexes for Cryo-EM studies

165 µM RecA_Pa_ was incubated over the weekend on ice with 13 µM 72mer oligo(dT) ssDNA and 1 mM ATPγS to induce RecA_Pa_ oligomerization on ssDNA. The sample was either diluted three times in 10 mM HEPES, pH 7.1, 150 mM NaCl (*RecA_Pa_\** sample) or supplemented with 53 µM LexA_Pa_S125A and incubated 2h at 4 °C (*RecA_Pa_*-LexA_Pa_S125A* sample). Samples underwent protein crosslinking by adding 2.5 mM disuccinimidyl suberate (DSS; 5% v/v DMSO) and incubating overnight at 4 °C under gentle agitation. Reactions were quenched by adding 100 mM Tris (pH 7.0) for 2h at room temperature. Protein pellet was removed by centrifugation before loading the mixture on a Superdex 200 10/300 GL size-exclusion chromatography column (Cytiva), pre-equilibrated with 20 mM Tris-HCl, pH 7.5, 150 mM NaCl.

As revealed by electron microscopy preliminary observation of the different samples recovered, the helical nucleoprotein complexes were eluted with the void volume of the column.

Samples were concentrated by Vivaspin centrifugal devices (MWCO 50 kDa; Sartorius) before deposition on grids for cryogenic electron microscopy (Cryo-EM).

### Cryo-EM data collection

3 µL of freshly purified RecA_Pa_* complex (2.3 mg/mL) were applied to a glow discharged Quantifoil R 1.2/1.3 Cu300 holey carbon grid. Excess sample was blotted away, and the grid was plunge-frozen into liquid ethane using a Mark IV Vitrobot (1.0 s blot time, 10°C, 100% humidity) at the Florence Center for Electron Nanoscopy (Dept. of Chemistry, University of Florence, Italy). The grids were imaged on the 300 kV Titan Krios microscope (Thermo Fisher Scientific) of the CM01 facility of the ESRF (Kandiah *et al*, 2019) with a K2 direct electron detector camera (Gatan, USA) operated in counting mode and at a pixel size of 0.827 Å per pixel. A total of 8711 movies were collected with 50 frames each, a fractional exposure of 0.98 e^-^/Å^2^ per frame and using a defocus range from −0.8 to −2.0 µm.

3 μL of RecA_Pa_*-LexA_Pa_S125A were applied to a glow-discharged Quantifoil R 1.2/1.3 copper 300-mesh holey carbon grids. The grid was blotted and plunge-frozen as reported above. The grids were imaged on the 300 kV Titan Krios microscope (Thermo Fisher Scientific) of the CM01 facility of the ESRF (Kandiah *et al*, 2019) with a K3 direct electron camera (Gatan, USA) operated in counting mode and at a pixel size of 0.84 Å per pixel. A total of 7882 movies were collected with 54 frames each, a fractional exposure of 1.02 e^-^/Å^2^ per frame and using a defocus range from −1 to −2.0 μm in 0.2 μm steps. The exposure rate was 16.9 e^-^/pixel/sec for a total nominal exposure of 55.08 e^-^/Å^2^.

### Image processing and 3D reconstruction

For both datasets, motion correction was performed by Motioncor2 (Zheng *et al*, 2017) and parameters of the contrast transfer function (CTF) were estimated by Gctf (Zhang, 2016). For the RecA_Pa_* dataset, 6543 micrographs were selected for the analysis. A small set of filaments was manually traced from a subset of micrographs to obtain initial 2D class averages for use as templates for reference-based autopicking in RELION 3.1.1 (Zivanov *et al*, 2018). 609530 segments were automatically picked and extracted to a box size of 384 × 384 pixels with an overlap of 85% and imported into CryoSPARC v4.2.1 (Punjani *et al*, 2017). Following 2D classification, 202842 segments were selected and used for 3D refinement, using helical parameters already reported for the RecA* homolog from *E. coli* as starting values (i.e., helical twist = 59° and rise = 15.5 Å; Gao *et al*, 2023). Particles were further subjected to local and global CTF refinement yielding a consensus map at 4.2 Å overall resolution (Supplementary Fig 2) with final helical parameters as reported in Supplementary Table 2.

For RecA_Pa_*-LexA_Pa_S125A initial attempts of automatic picking failed. Therefore 104800 tubes were manually picked and extracted to a box size of 384 × 384 pixels with an overlap of 85% in RELION 4.0.0 (Kimanius *et al*, 2021). After several rounds of 2D classification, 561719 particles were used as input for the generation of a 3D initial model with C1 symmetry using a spherical mask of 350 Å, in RELION 4.0.0. After import into CryoSPARC v4.2.1 (Punjani *et al*, 2017) and heterogeneous refinement, 438072 particles were selected for a further round of homogenous refinement. Following two rounds of 3D classification, first using a spherical mask of radius 50 Å centered on LexA density, and then a structure-based mask encompassing the LexA_Pa_ density, were used to select 164165 particles, corresponding to the classes presenting additional density in the RecA_Pa_* groove which we ascribed to LexA_Pa_. After local and global CTF refinement, homogenous refinement led to a consensus map at 3.4 Å overall resolution (Supplementary Fig. 3).

Local amplitude scaling was performed using the model-free implementation of local sharpening with reference profiles in LocScale2 (Jakobi *et al*, 2017; Bharadwaj & Jakobi, 2022) with a cubic averaging window of 25 Å edge length and starting from the unfiltered half maps. The locally scaled map was used for display purposes (Fig. 2 A-B, Fig. 3 A-B, Supplementary Fig. 2 and 3); atomic model refinement and model-map FSC calculations were done using the original half maps.

### Model building, refinement and structural analysis

A homology model for the atomic structure of monomeric RecA_Pa_ was generated by SwissModel (Waterhouse *et al*, 2018) using PDB 2REB (monomeric *E. coli* RecA) as a template. The model was fitted into a zone corresponding to a single RecA_Pa_ monomer in the cryo-EM map. Then the full oligomer was reconstructed by applying the helical symmetry parameters using UCSF Chimera. The ssDNA poly(dT) chain, ATPγS and Mg^2+^ ions were built and fitted using Coot (Emsley *et al*, 2010). The resulting model was refined by iterative cycles of automated real space refinement in Phenix (Afonine *et al*, 2012). For RecA_Pa_*-LexA_Pa_S125A, our structure of RecA_Pa_* was used as the starting model. LexA_Pa_^CTD^ in the cleavable conformation (i.e., with the cleavable loop closed) was modeled by Phyre2 web server in the “one-to-one threading” mode (Kelley *et al*, 2015), using LexA ^CTD^ from PDB 8GMS as the template. The closed cleavable loop was then grafted on chain A of LexA_Pa_^CTD^ G91D X-ray structure and the S125A mutation was introduced by Pymol v2.3.5. RecA_Pa_* and the model of LexA_Pa_^CTD^ dimer were fitted into the respective densities in the map and refined by automatic and manual real-space refinement methods using Phenix in the default mode (Afonine *et al*, 2018) and Coot (Emsley *et al*, 2010), respectively. Analysis of protein-protein and protein-ligand interactions was performed by PDBePISA (https://www.ebi.ac.uk/pdbe/pisa/; Krissinel & Henrick, 2007) and PLIP (https://plip-tool.biotec.tu-dresden.de/plip-web/plip/index; Adasme *et al*, 2021).

### Fluorescence polarization-based studies

Fluorescence polarization (FP) was used as the biophysical readout to observe and quantify the binding of RecA_Pa_ to ssDNA and LexA_Pa_ to RecA_Pa_*.

To determine the apparent affinity of RecA_Pa_ for ssDNA and ATPγS, a 5’-Carboxyfluoresceinated 32- mer oligonucleotide (FAM-32mer; Supplementary Table 3) was used as “scaffold” (Lee *et al*, 2007; Cory *et al*, 2022).

In the former experiment, 10 nM FAM-32mer ssDNA was incubated with different concentrations of RecA_Pa_ and an excess of ATPγS (1 mM) for 30 min at 37 °C before reading the FP signal. FP data measured without RecA_Pa_ and at 17 µM RecA_Pa_ were considered as “0% oligomerization” and “100% oligomerization”, respectively, and used to normalize all the collected data, thus deriving the fraction of RecA_Pa_-bound ssDNA in each sample. The RecA_Pa_-bound fraction (*F_B_*) of FAM-32mer ssDNA was plotted against RecA_Pa_ concentration and experimental data were best-fitted in GraphPad Prism 8 by a Hill equation (eq. 1, where *h* is the Hill coefficient; Stefan & Le Novère, 2013; Jarmoskaite *et al*, 2020; Gesztelyi *et al*, 2012).

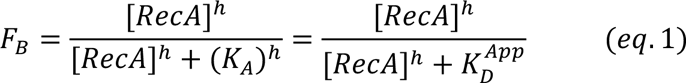

Conversely, to estimate RecA_Pa_ apparent affinity for ATPγS, 10 nM FAM-32mer ssDNA and 1 µM RecA_Pa_ were incubated with different concentrations of ATPγS for 30 min at 37 °C before reading the FP signal. FP data measured without ATPγS and at 10 mM ATPγS were considered as “0% oligomerization” and “100% oligomerization”, respectively, and used to normalize all the collected data. The RecA_Pa_-bound fraction of FAM-32mer ssDNA was plotted against ATPγS concentration and experimental data were best-fitted in GraphPad Prism 8 by a Hill equation (eq. 2).

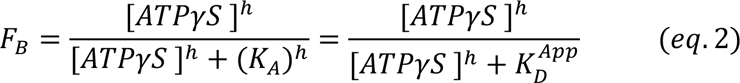

To determine the apparent affinity of LexA_Pa_ to RecA_Pa_*, FlAsH-LexA_Pa_^CTD^S125A was used as the fluorescent probe at a fixed concentration of 0.1 µM. RecA_Pa_ was pre-activated with SKBT25-18mer ssDNA and ATPγS and then added at different concentrations. Following a 30 min incubation at 37°C, the FP signal was measured. FP data measured without RecA_Pa_* (0% binding) and at 10 µM RecA_Pa_* (100% binding) were used to normalize all the data and obtain the RecA_Pa_*-bound fraction of FlAsH-LexA_Pa_^CTD^ S125A. Normalized data were best-fitted in GraphPad Prism 8 by a single binding site model.

---

**Supplementary Fig. 1:**
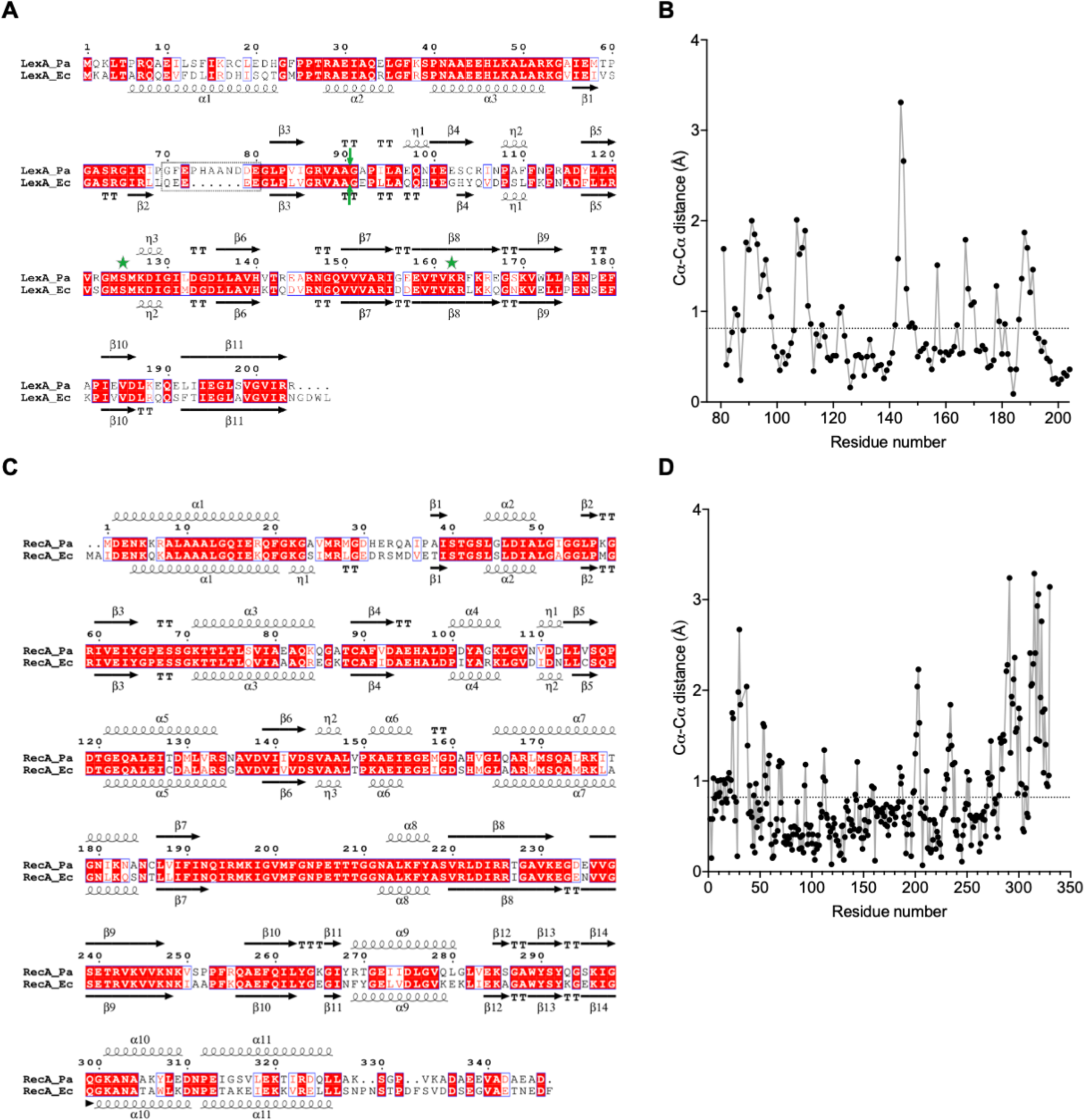
Sequence and structure comparison of LexA and RecA from *P. aeruginosa* and *E. coli*. (**A**) Sequence alignment of LexA_Pa_ and LexA_Ec_. Secondary structures are indicated, as observable in PDB 8B0V and PDB 3JSO, respectively. Residues of the catalytic Ser/Lys dyad are indicated by green stars, while the cleavage site is indicated by green arrows. (**B**) Structural comparison between LexA_Pa_^CTD^ (PDB 8B0V) and LexA_Ec_^CTD^ (PDB 1JHF) by Gesamt. The average Cα-Cα distance is shown as a dotted line. (**C**) Sequence alignment of RecA_Pa_ and RecA_Ec_. Secondary structures are indicated, as observable in PDB 8S70 and PDB 7JY6, respectively. (**D**) Structural comparison between RecA_Pa_ (PDB 8S70) and RecA_Ec_ (PDB 7JY6) by Gesamt. The average Cα-Cα distance is shown as a dotted line.

**Supplementary Fig. 2:**
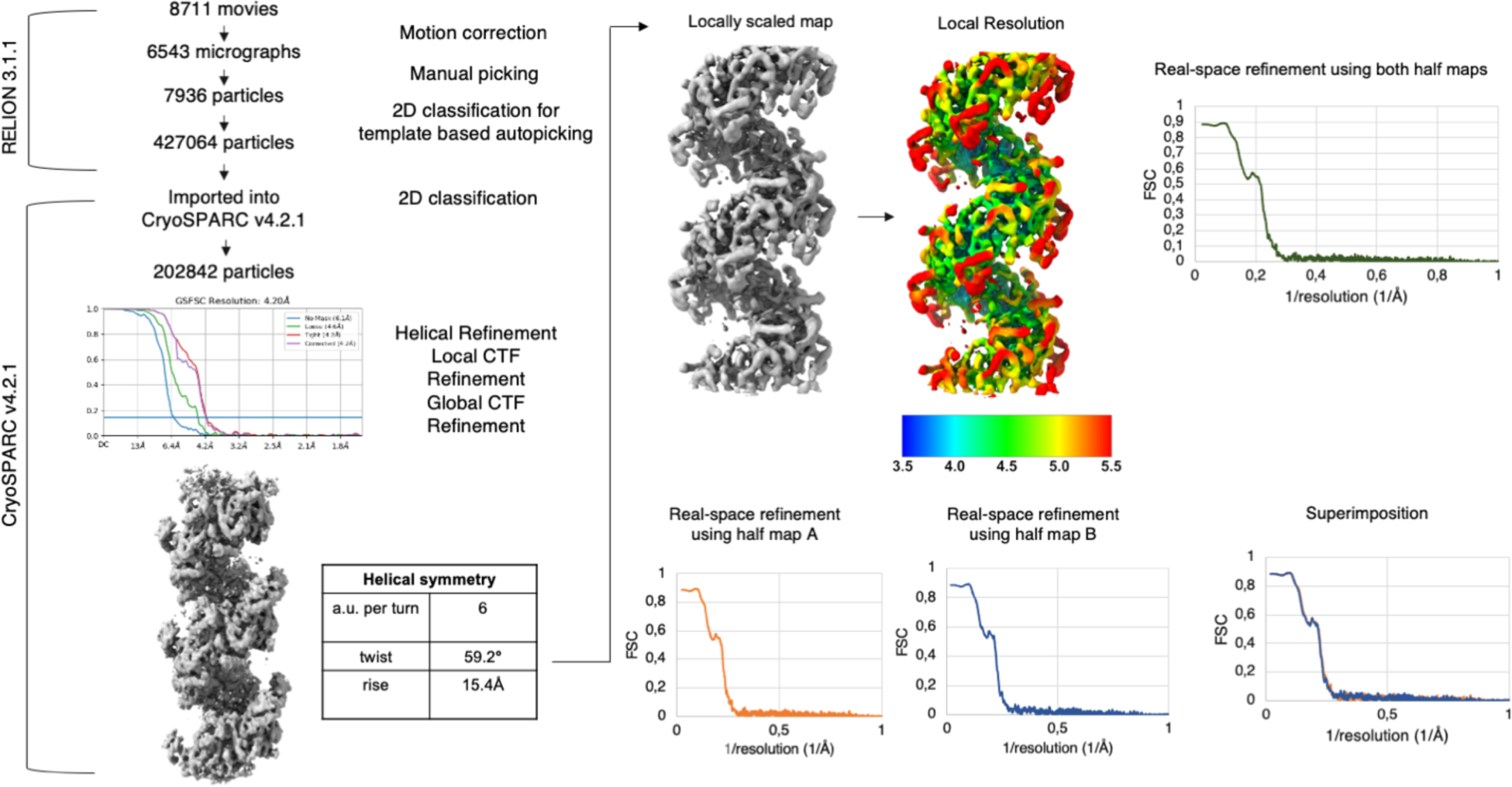
RecA_Pa_* Cryo-EM analysis pipeline. Details of the data processing from movie alignment to final helical refinement are shown.

**Supplementary Fig. 3.:**
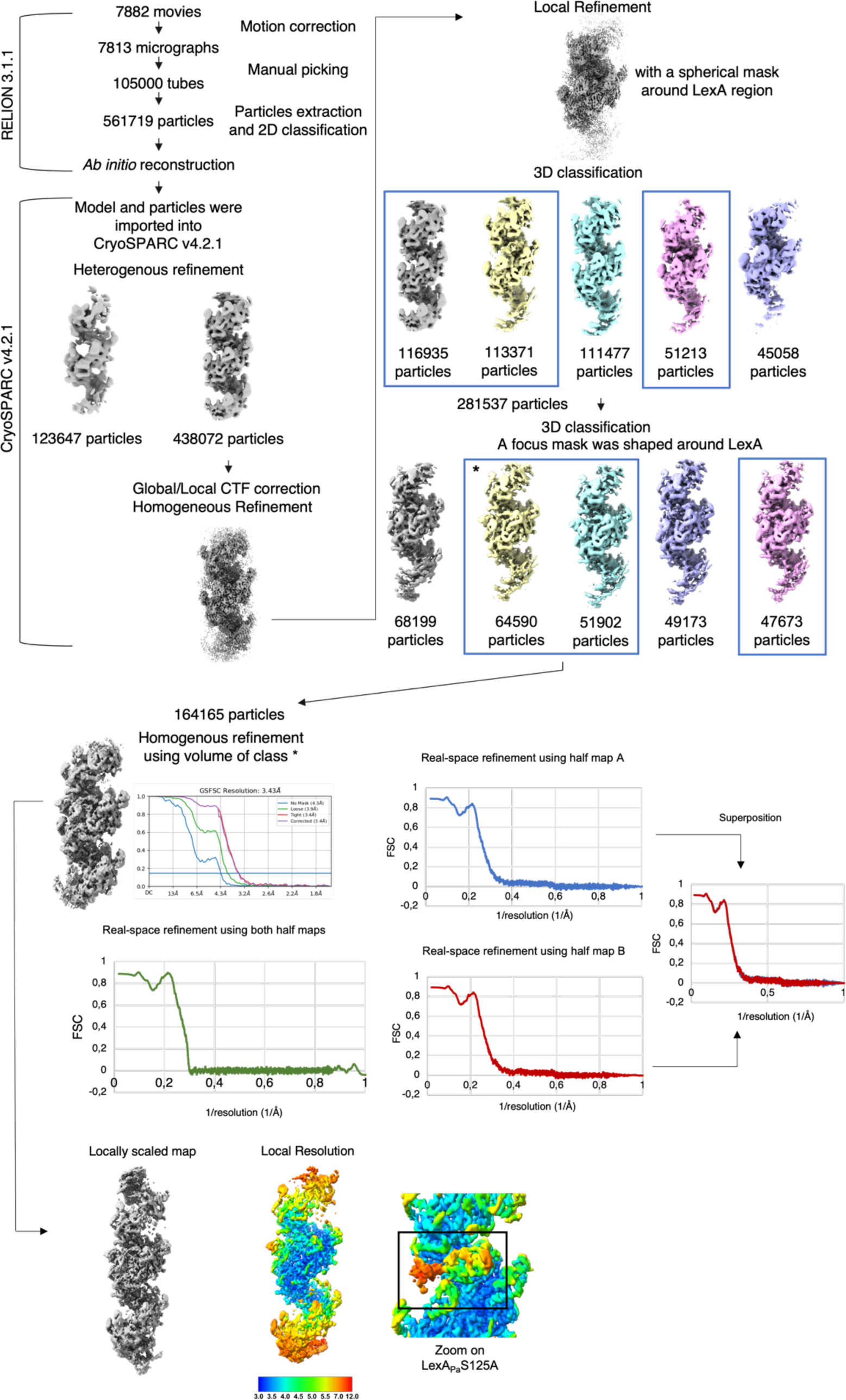
RecA_Pa_*-LexA_Pa_S125A Cryo-EM analysis pipeline. Details of data processing, from movie alignment to final refinement are shown.

**Supplementary Fig. 4.:**
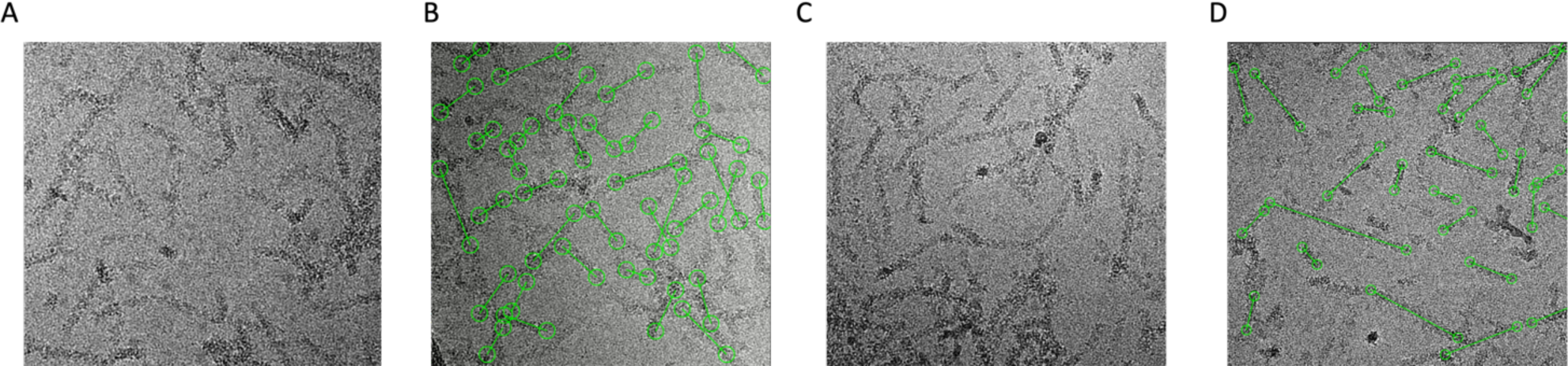
Representative micrographs (A/C) and manually picked ones (B/D) of RecAPa* and the complex of RecAPa* and LexAPa S125A, respectively.

**Supplementary Table 1:**
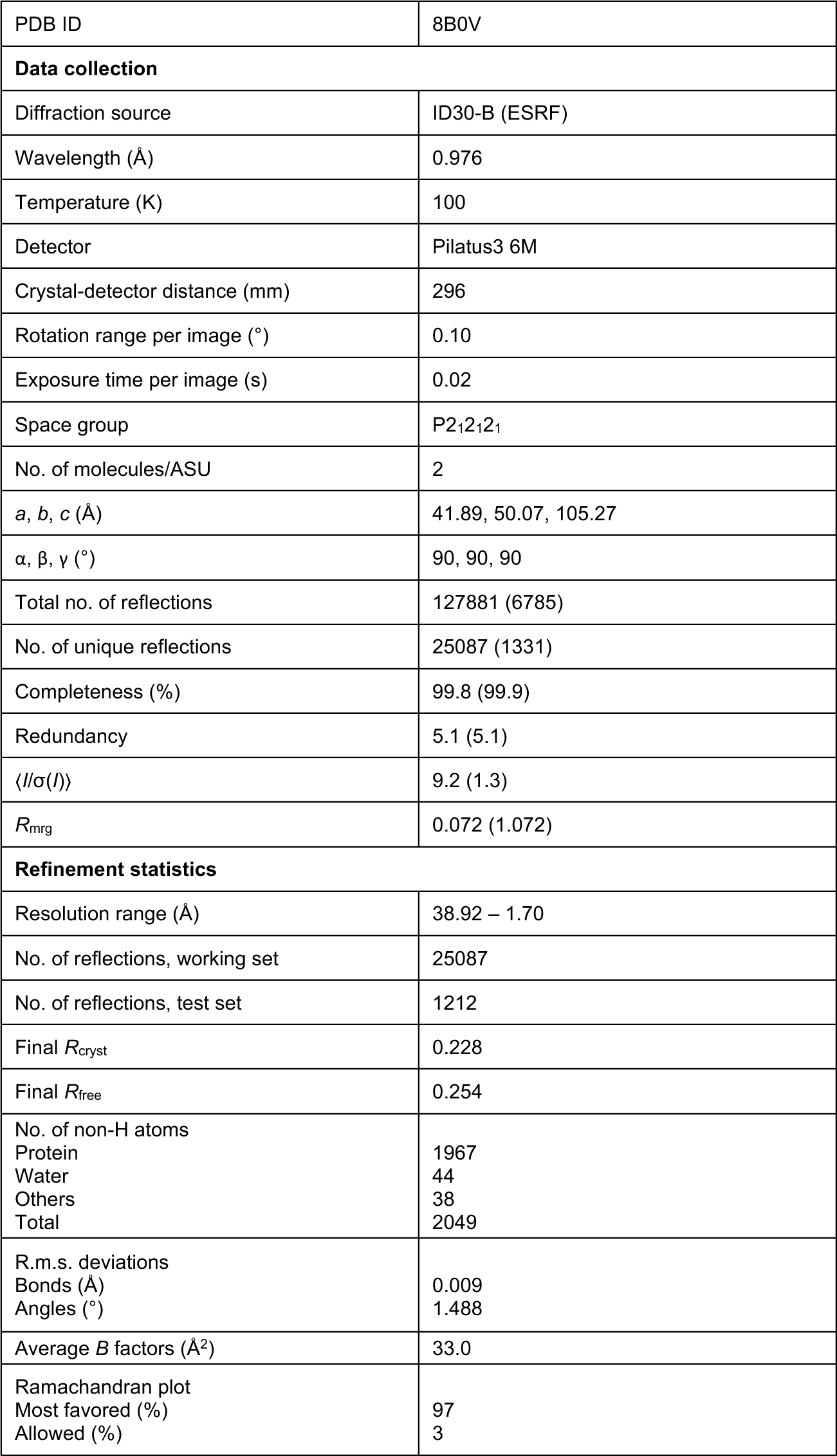
X-ray data collection and refinement statistics.

|  |  |
| --- | --- |
| PDB ID | 8B0V |
| **Data collection** |  |
| Diffraction source | ID30-B (ESRF) |
| Wavelength (Å) | 0.976 |
| Temperature (K) | 100 |
| Detector | Pilatus3 6M |
| Crystal-detector distance (mm) | 296 |
| Rotation range per image (°) | 0.10 |
| Exposure time per image (s) | 0.02 |
| Space group | P2<sub>1</sub>2<sub>1</sub>2<sub>1</sub> |
| No. of molecules/ASU | 2 |
| *a*, *b*, *c* (Å) | 41.89, 50.07, 105.27 |
| $\alpha$, $\beta$, $\gamma$ (°) | 90, 90, 90 |
| Total no. of reflections | 127881 (6785) |
| No. of unique reflections | 25087 (1331) |
| Completeness (%) | 99.8 (99.9) |
| Redundancy | 5.1 (5.1) |
| $\langle I/\sigma(I) \rangle$ | 9.2 (1.3) |
| *R*<sub>merge</sub> | 0.072 (1.072) |
| **Refinement statistics** |  |
| Resolution range (Å) | 38.92 – 1.70 |
| No. of reflections, working set | 25087 |
| No. of reflections, test set | 1212 |
| Final *R*<sub>cryst</sub> | 0.228 |
| Final *R*<sub>free</sub> | 0.254 |
| No. of non-H atoms |  |
| Protein | 1967 |
| Water | 44 |
| Others | 38 |
| Total | 2049 |
| R.m.s. deviations |  |
| Bonds (Å) | 0.009 |
| Angles (°) | 1.488 |
| Average *B* factors (Å<sup>2</sup>) | 33.0 |
| Ramachandran plot |  |
| Most favored (%) | 97 |
| Allowed (%) | 3 |

**Supplementary Table 2:**
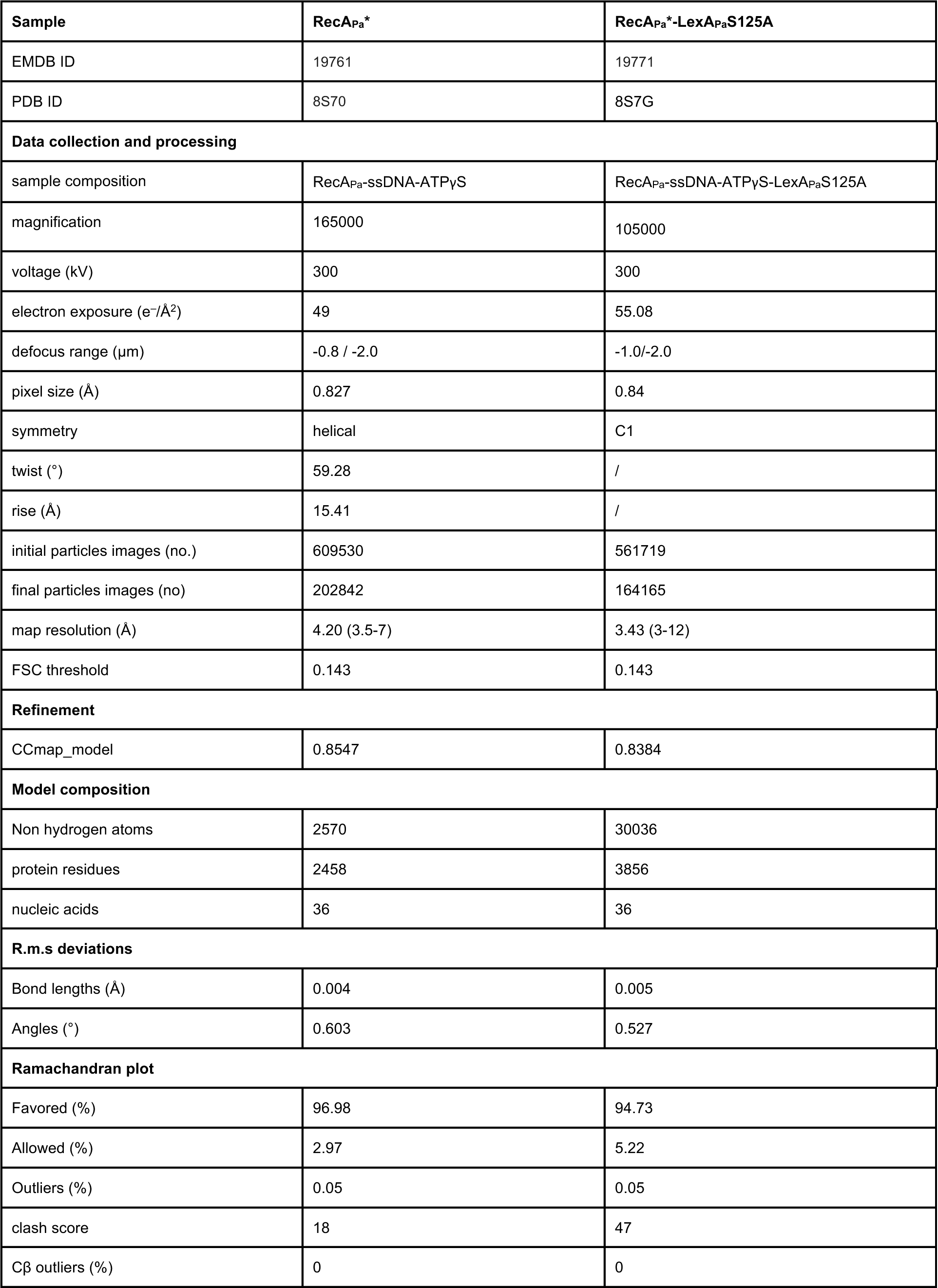
**Cryo-EM data collection, processing, and structure refinement statistics**.

**Supplementary Table 3:**
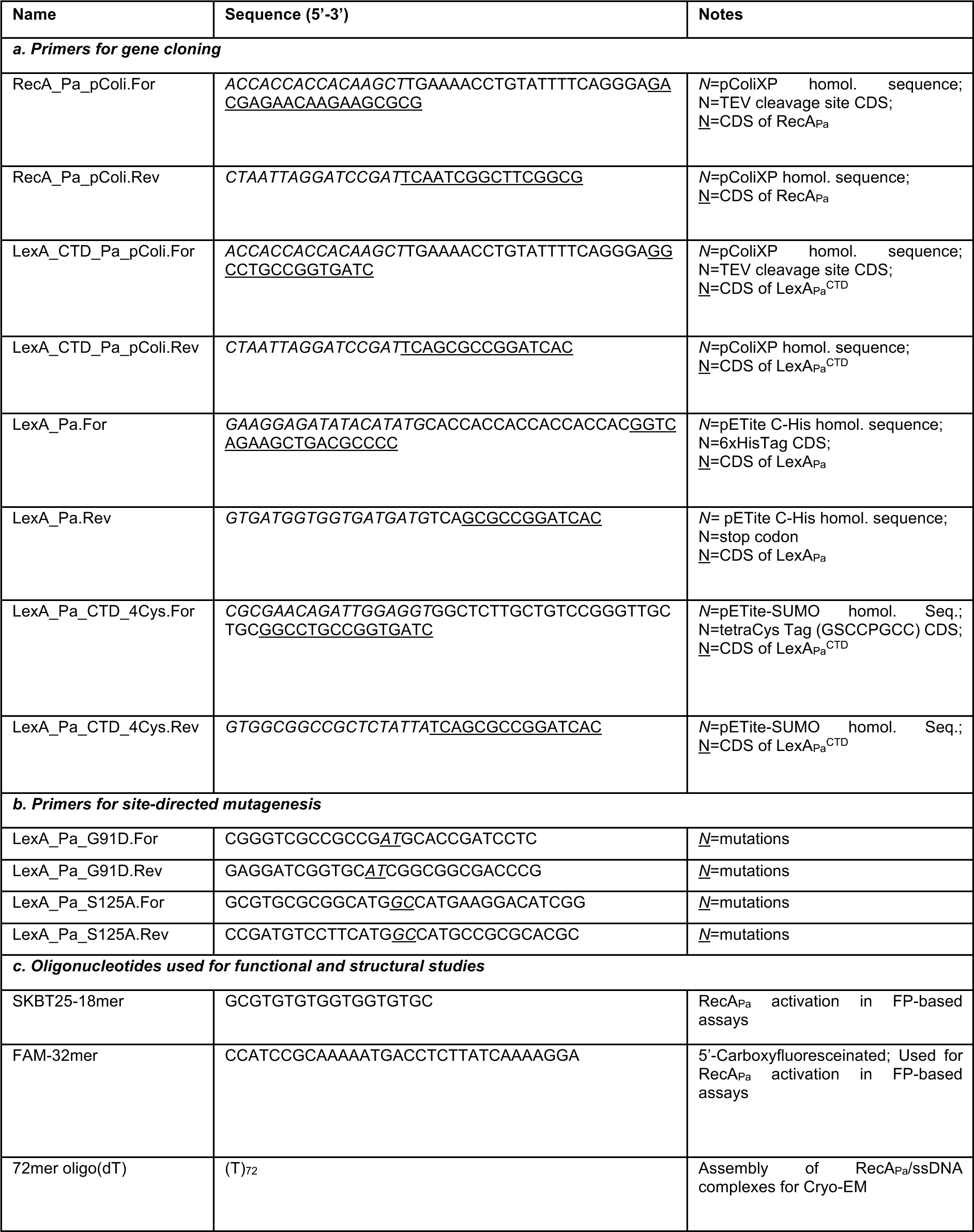
**Oligonucleotides**.

## Notes

### Competing Interest Statement

The authors have declared no competing interest.

